# Dynamic acoustic-to-categorical representations of phonemes and prosody along ventral and dorsal speech streams

**DOI:** 10.1101/2025.01.24.634030

**Authors:** Seung-Cheol Baek, Seung-Goo Kim, Burkhard Maess, Maren Grigutsch, Daniela Sammler

## Abstract

Phonemes and prosodic contours are fundamental elements of speech used to convey complementary meanings. Perceiving these elements requires mapping variable acoustic cues onto discrete categories along ventral and dorsal speech streams. While traditional models make clear predictions, exactly where and when this acoustic-to-categorical mapping occurs remains unclear. Using magnetoencephalography and behavioural psychophysics, combined with time-resolved representational similarity and multivariate transfer entropy analyses, we show how phonemes and prosody propagate along the dual streams and how their categorical representations are gradually formed. Contrary to theoretical predictions, acoustic and categorical representations occur in parallel, rather than serially, across time and space for both elements. Moreover, prosody categories extend further along both streams than phoneme categories, with differently weighted contributions of posterior temporal areas. These results highlight a shared principle of parallel acoustic and categorical processing, yet partially distinct abstraction mechanisms for phonemes and prosody, key to access the multilayered meaning of speech.

## Introduction

Speech conveys meaning at multiple levels. The propositional meaning coded by words— composed of phonemes—is often complemented by speaker meaning, that is, a speaker’s communicative intentions conveyed by the tone of voice—the prosody—of an utterance^1^. A crucial first step in fully comprehending both levels is to perceive the fundamental elements that underlie these meanings from the myriad variable acoustic cues in the dynamic speech stream. Transient spectrotemporal cues code phonemes^2^, which are combined to form words, while characteristic patterns of fundamental frequency (*F*_0_) over time form prosodic pitch contours (henceforth, prosody) that mark, for example, whether a speaker intends to make a statement or ask a question^3^. Although we often effortlessly perceive discrete categories of phonemes and prosody from the relevant acoustic cues^4,5^, the neural substrate underlying this acoustic-to-categorical mapping remains poorly understood.

Speech processing is known to involve distributed frontotemporal areas, organised into ventral and dorsal streams^6–8^. Classical hierarchical processing models propose that, within this dual-stream architecture, continuously variable acoustic signals are serially transformed into more abstract linguistic representations across both time and space^6,9–11^. That is, speech is first acoustically analysed in sensory areas and then represented as discrete categories of linguistic elements in higher-order regions, including lateral and anterior temporal^12–14^, motor^15,16^ or inferior frontal regions^17,18^.

However, this simple serial hierarchy has recently been challenged in several ways— for example, by studies showing categorical representations in the primary auditory cortex^19–21^, or direct encoding of abstract linguistic information in the superior temporal cortex, bypassing sensory areas^22^. Although evidence from functional neuroimaging or electrophysiology has provided some indications of where and when acoustic-phonetic^23–26^ or categorical^17,21,27–29^ features of speech are represented in the brain, studies have often been unable to effectively dissociate these two representations, leaving it elusive how the acoustic-to-categorical mapping actually occurs. In particular, the categorical encoding reported in prior studies, especially those using naturalistic stimuli, often lacks perceptual relevance^30^, as it is not directly linked to actual categorical decisions^21,27,28^, and may instead reflect acoustic confounds^31^ driven by the intrinsically high correlation between acoustic and categorical representations^2,32^. Moreover, the limited spatial or temporal resolution of non-invasive methods and the restricted coverage of intracranial recordings hamper understanding the precise spatiotemporal profiles of acoustic and categorical encoding at the network level. Specifying these profiles has major implications for understanding how the brain forms abstract linguistic representations from a dynamic speech signal.

Notably, almost all studies seeking to uncover the acoustic-to-categorical mapping in speech have focused on phonemes^15,17,20,29,33^, leaving one important aspect of speech largely unexplored: speech prosody. Compared to phonemes, prosody is not only less well established in its categorical perception but is also characterised by distinct acoustic cues; whether it engages similar neural processing is therefore still an open question. While traditional views posit a shared acoustic analysis^11^, phonemes and prosody are often claimed to tap into different hemispheric preferences^7,8,13^ or involve segregated encoding across the temporal cortex^22,25^. Determining the extent to which the neural infrastructure supporting phonemic and prosodic processing is shared across the dual-stream networks is critical for understanding the general computational principles underlying the perception of the fundamental building blocks of speech at different timescales.

To address these gaps, we acquired magnetoencephalography (MEG) data from 29 participants during a psychophysical experiment with single words varying in word-initial phoneme and prosodic pitch contour. Building on participants’ behaviour and MEG recordings, we combined time-resolved representational similarity analysis^34,35^ (RSA) with multivariate transfer entropy^36,37^ (mTE) analysis to reliably trace the neural representations of phonemes and prosody at different levels of abstraction (i.e., acoustic and categorical) across space and time, as well as their information flow between the involved regions.

Here, we show the abstraction and propagation of both phonemes and prosody along the dual speech streams. Notably, our results indicate that the acoustic-to-categorical transformation is not exactly serial as proposed by classical models^6,9–11^. Instead, the two levels of abstraction are processed largely in parallel across time and space. The convergence of these findings for both linguistic elements points to a shared principle of representational abstraction. Marked differences are the spatially broader categorical representations of prosody compared to phonemes along both steams, and the differently weighted contributions of posterior temporal areas. This suggests partially distinct abstraction mechanisms that warrant confirmation in future studies. Together, our findings demonstrate that the brain abstracts both phonemes and prosody through parallel processing of acoustic and categorical information within the dual-stream networks, while flexibly adapting regional involvement and information flow to meet the demands posed by the respective cues and linguistic functions.

## Results

To track how phonemic and prosodic representations dynamically evolve in the brain, we created a stimulus set of 7,442 mono-syllabic words that varied in 61 evenly spaced steps along two orthogonal dimensions (middle panel of Fig. 1a): (i) the voice onset time (VOT) of the word-initial phoneme shifting from /b/ to /p/, and (ii) the pitch contour of the word shifting from statement (falling) to question (rising). Stimuli were generated via audio morphing of natural recordings of the German words, “Bar” (Engl., “pub”) and “Paar” (Engl., “pair”), uttered as both statement and question by two German native speakers (one female). A subset of 25 stimuli per speaker, that is, 5 x 5 morph levels out of the 61 x 61 steps, were selected individually for each of our 29 participants (Fig. 1a). The selection was based on participants’ performance in pre-tests (see Supplementary Methods), which served to centre stimuli on the individual phonemic and prosodic category boundaries, and to match the difficulty of categorising the phonemes and prosody, for both speakers.

**Fig. 1.**
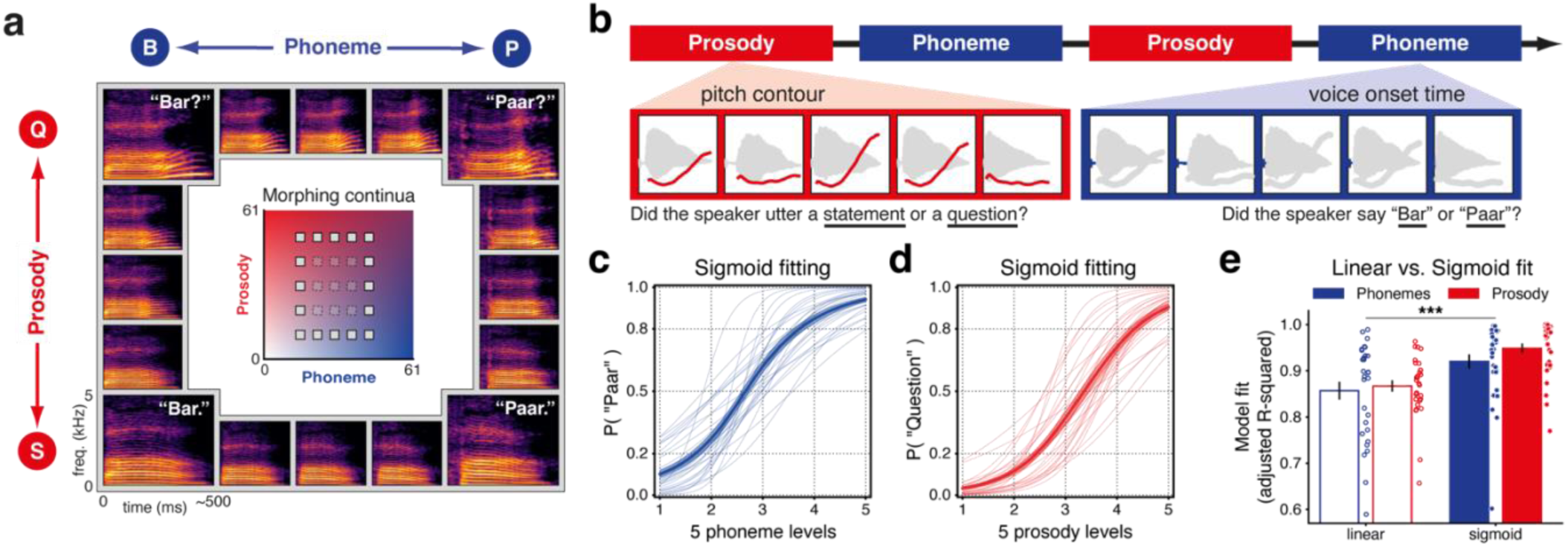
Phoneme and prosody identification tasks and behavioural results. **a**, Auditory stimuli consisted of single words that varied in word-initial phonemes (horizontal axis) and prosodic pitch contours (vertical axis). The stimuli were generated via audio morphing applied to natural recordings of two German words, “Bar” (B; Engl., “pub”) and “Paar” (P; Engl., “pair”), uttered as both statement (S) and question (Q). Morphing resulted in a stimulus space of 61 x 61-step orthogonal continua (middle panel) with gradual transitions from /b/ to /p/ and from statement to question. 5 x 5 levels in this space, exemplified by the small squares, were selected for each participant (see also Fig. 2d). Stimulus examples are depicted as spectrograms in the surrounding panels (corresponding to the white squares in the stimulus space). **b**, Schematic of one out of six experimental runs. Each run contained two phoneme and two prosody blocks in an alternating order, in which participants were asked to identify the word (“Bar”/“Paar”) or the prosody (“statement”/“question”), respectively. **c**,**d**, Sigmoid functions fitted to behavioural responses of “Paar” across five phoneme levels (**c**) and of “question” across five prosody levels (**d**). Shaded areas indicate the standard error of the mean (SEM). Thin lines represent individual participants. **e**, Comparisons of linear and sigmoid fits (adjusted R-squared) across the two identification tasks. Irrespective of the tasks, behavioural responses across five morph levels better followed a sigmoid function than a linear function (repeated measures analysis of variance, main effect of function, *p* < 0.001; ***), confirming that both phonemes and prosody are perceived categorically. Error bars denote SEM. Linear (empty) and sigmoid (filled) fits for each participant are depicted as circles.

During the MEG recordings, participants were asked in two-alternative forced-choice manner whether the speaker said “Bar” or “Paar,” or uttered “statement” or “question,” in alternating phoneme and prosody blocks presenting the 25 distinct stimuli of one speaker (Fig. 1b). The comparison of the slopes of the psychometric functions fitted to the proportions of “Paar” or “question” responses per speaker and morph level along the respective dimension confirmed that stimuli were matched in difficulty as intended^38^: Slopes differed neither between tasks (*F*_1,28_ = 0.203, *p* > 0.05, 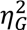 = 0.002) nor speakers (*F*_1,28_ = 1.105, *p* > 0.05, 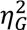 = 0.012), and no significant interaction was observed (*F*_1,28_ = 1.385, *p* > 0.05, 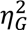 = 0.009). Moreover, the point of subjective equality (PSE), although different between tasks (*F*_1,28_ = 39.852, *p* < 0.001, 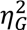 = 0.266), did not differ between speakers (*F*_1,28_ = 0.766, *p* > 0.05, 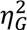 = 0.000; no task x speaker interaction: *F*_1,28_ = 0.485, *p* > 0.05, 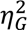 = 0.004), allowing us to pool data across speakers in the remaining analyses.

### Categorical perception of phonemes and prosody

We first confirmed that the stimuli were perceived categorically, both in terms of phonemes and prosody. To do so, we compared the goodness of fit between linear and psychometric functions fitted to the behavioural responses (Fig. 1c-e). The psychometric function provided a significantly better fit than the linear function (*F*_1,28_ = 109.733, *p* < 0.001, 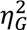 = 0.173; Fig. 1e), similarly in both tasks (no main effect of task: *F*_1,28_ = 1.182, *p* > 0.05, 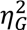 = 0.014; no task x function interaction: *F*_1,28_ = 1.012, *p* > 0.05, 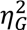 = 0.003). These findings indicate that not only phonemes but also prosody are perceived categorically; that is, continuous acoustic cues are mapped onto discrete categorical representations, similarly along both of our stimulus dimensions.

### Phonemic and prosodic representations in ventral and dorsal stream regions

We then explored where and when phonemic and prosodic information is generally represented in the brain by applying time-resolved RSA to MEG source activity pooled across both tasks^34,35^. In this analysis, time-varying dissimilarities of neural activity patterns evoked by the five phoneme or prosody levels were compared with modelled dissimilarities of both stimulus acoustics and perceived categories (Fig. 2).

**Fig. 2.**
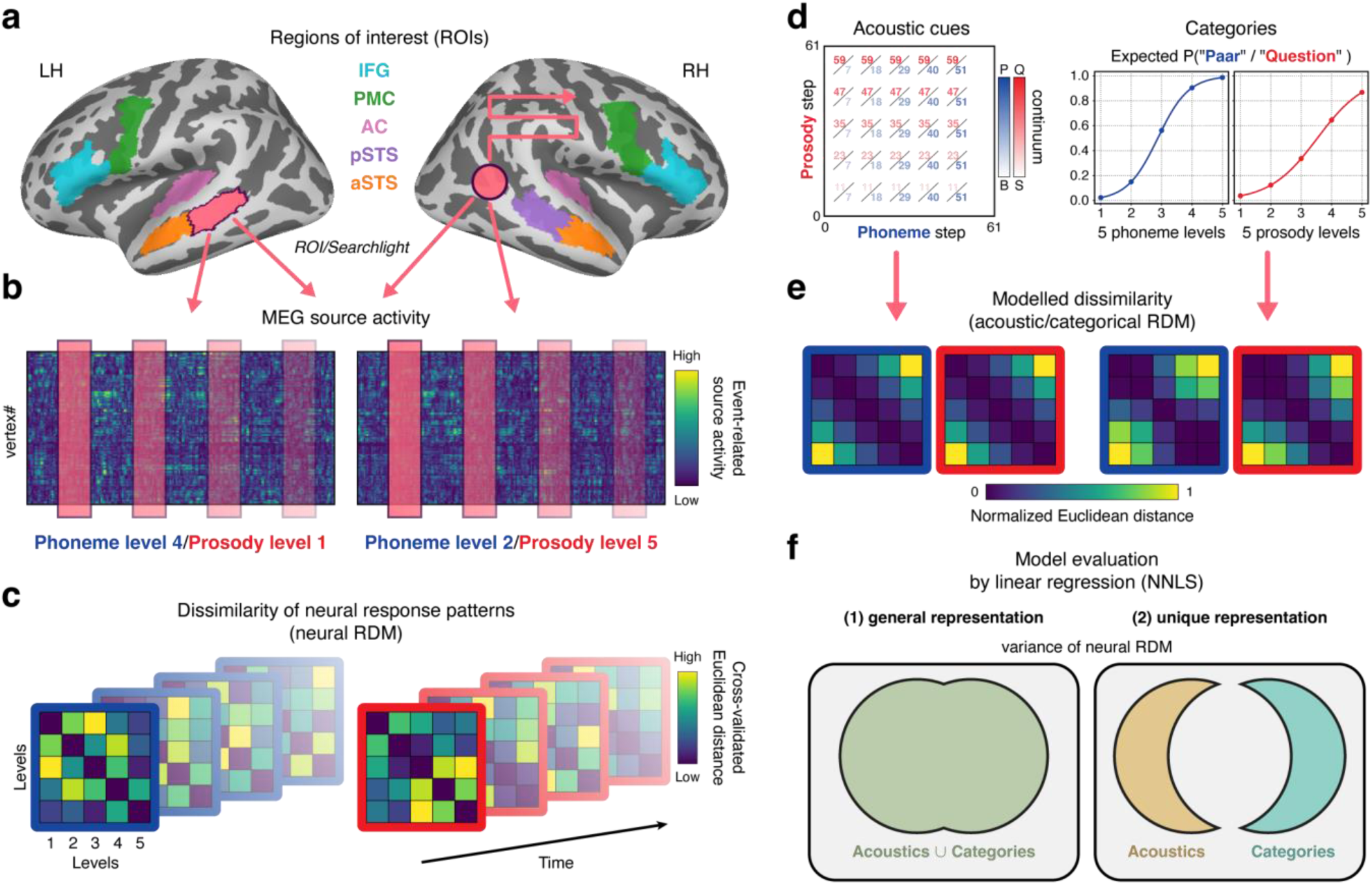
Workflow of time-resolved representational similarity analysis (RSA). **a**, Multivariate magnetoencephalography (MEG) source activity evoked by the five phoneme or prosody levels was extracted from 10 predefined regions of interest (ROIs) across the left (LH) and right (RH) hemispheres for each participant. The ROI-based approach was complemented by a whole-brain searchlight analysis (searchlight radius = 10 mm; light red circle). IFG, inferior frontal gyrus; PMC, premotor cortex; AC, auditory cortex; p/aSTS, posterior/anterior superior temporal sulcus. **b**, For each ROI or searchlight location, cross-validated squared Euclidean distance (over six experimental runs) was calculated between the local neural response patterns to the five morph levels in 24-ms sliding time windows (step size: ROI = 4 ms, searchlight = 12 ms). **c**, This yielded a time series of 5 x 5 neural representational dissimilarity matrices (RDMs) for each participant, separately for phonemes (blue) and prosody (red). **d**, Neural representational characteristics were probed in terms of both stimulus acoustics and perceived categories. Stimulus acoustics (left) were coded as the original morph step of the stimuli on the 61-step phoneme (B, “Bar”; P, “Paar”) or prosody (S, “statement”; Q, “question”) continua. Perceived categories (right) were soft-coded as the expected proportion of “Paar” or “question” responses at each of the five levels, derived from the sigmoid functions fitted to individual behaviour. **e**, Stimulus acoustics and perceived categories across the five levels were compared separately using squared Euclidean distance, normalised between 0 and 1. This resulted in one acoustic and one categorical RDM, each with a size of 5 x 5, per participant, for phonemes and for prosody. **f**, To assess general representations of phonemes and prosody for each participant (left), each time step of the corresponding neural RDMs in a given region was predicted by a linear combination of acoustic and categorial RDMs using non-negative least squares (NNLS) fitting. Explained variance was further partitioned to examine unique acoustic and categorical representations (right).

As illustrated in Fig. 2a, neural dissimilarities were quantified in 10 predefined regions of interest (ROIs) across the two hemispheres, including the inferior frontal gyri (IFG) and premotor cortices (PMC) in the dorsal stream, as well as the auditory cortices (AC) and the posterior (pSTS) and anterior superior temporal sulci (aSTS) in the ventral stream. In addition, we introduced a whole-brain searchlight approach to complement the ROI-based analysis. In each ROI or searchlight, we generated time series of 5 x 5 representational dissimilarity matrices (RDMs) based on the cross-validated squared Euclidean distance between the local multivariate response patterns to the five morph levels^39^, separately for phonemes and prosody, in 24-ms sliding time windows (Fig. 2b,c).

Model RDMs were built based on stimulus acoustics and perceived categories of the stimuli using squared Euclidean distance (Fig. 2d). Stimulus acoustics were coded as the original morph step of each stimulus on the 61-step phoneme or prosody continua.

Perceived categories were soft-coded as the expected proportion of “Paar” or “question” responses at each of the five morph levels derived from the psychometric functions fitted to individual behaviour. Based on these values, four model RDMs were constructed for each participant, coding the acoustic and categorical dissimilarity of all 5 x 5 combinations of morph levels, separately for phonemes and prosody (Fig 2e).

To assess neural representations of phonemes and prosody, the corresponding model RDMs were used to predict each time step of the neural RDMs in a given region. We first measured the variance explained by a linear combination of both acoustic and categorical RDMs over time to identify general representations, and later applied variance partitioning to tease apart acoustic and categorical representations (Fig. 2f). As a control, surrogate and noise ceiling analyses validated that the variability in the neural RDMs was largely accounted for by the combined acoustic and categorical RDMs: The observed explained variance reached the empirical noise ceiling in all ROIs, and no significant representations were found when the five morph levels were permuted (Supplementary Fig. 1).

Fig. 3a shows that phonemic information was represented exclusively in ventral stream ROIs in both hemispheres, while dorsal stream areas remained silent. Phonemic representations in all six temporal ROIs emerged as early as 70 ms after stimulus onset, that is, shortly after voice onset (26–62 ms), with the first peak occurring in the right AC at 108 ms post-onset, followed by peaks in the other regions, all aligned around 160 ms post-onset (Table 1). Additionally, the bilateral AC and pSTS exhibited sustained phonemic representations after 300 ms post-onset, with those in the bilateral AC persisting beyond stimulus offset (464–500 ms). Converging results were found in the whole-brain searchlight analysis (Fig. 3a, Supplementary Fig. 2a). Overall, these results indicate a high redundancy of phonemic representations across bilateral temporal areas and time, raising the question whether and how phonemic information propagates along the ventral stream.

**Fig. 3.**
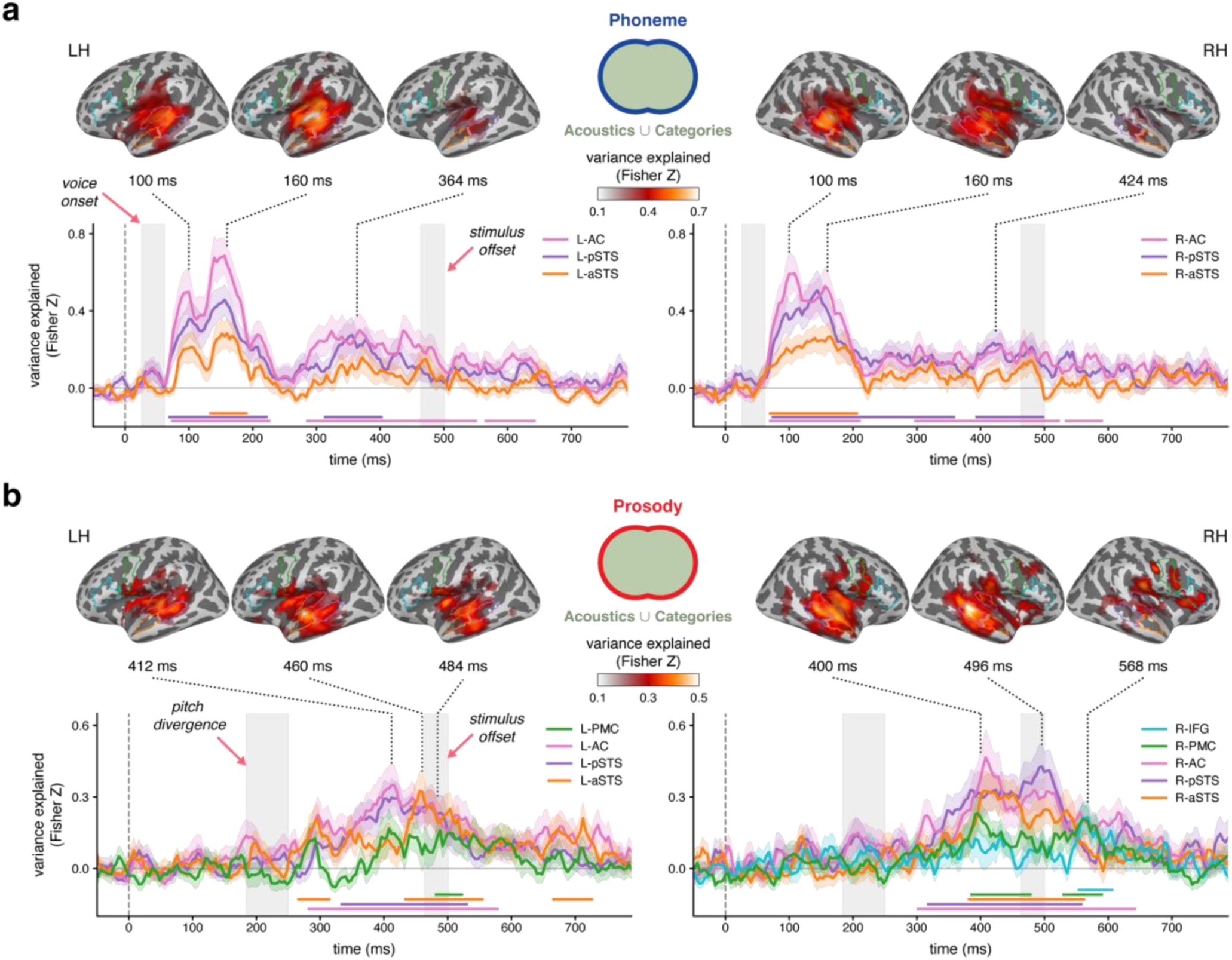
Phonemic and prosodic representations in ventral and dorsal stream regions. **a,b**, Time-varying variance (transformed into Fisher-Z scale and corrected by the average baseline from −200 to 0 ms around stimulus onset) explained by both acoustic and categorical RDMs in the left-(L-; left panel) and right-hemispheric (R-; right panel) ROIs that exhibit significant representations of phonemes (**a**) or prosody (**b**). Statistical significance was determined by cluster-based permutation tests (one-sided *t*_28_-test against zero, cluster-forming threshold (CFT) at *p* < 0.05, 100,000 permutations, family-wise error rate (FWER) = 0.05 across time points per ROI, ROI-wise false discovery rate (FDR) *q* < 0.05). ROI colour coding is the same as in Fig. 2a (IFG, inferior frontal gyrus; PMC, premotor cortex; AC, auditory cortex; p/aSTS, posterior/anterior superior temporal sulcus). Shaded areas indicate SEM. Horizontal lines above the x-axis show significant time intervals (ROI-wise cluster and peak statistics are summarised in Table 1). At selected time points around the peaks of the ROIs, cortical variance maps (Fisher-Z scale, baseline-corrected) of significant representations from the whole-brain searchlight analysis are shown at the top (cluster-based permutation tests, one-sided *t*_28_-test against zero, CFT at *p* < 0.05, 10,000 permutations, FWER = 0.05 across vertices and time points) for the left (LH) and right (RH) hemispheres, with the ROIs outlined in the corresponding colours for comparison. **a**, Phonemic representations emerged after voice onset (left grey rectangle) across bilateral temporal areas and persisted beyond stimulus offset (right grey rectangle), particularly in posterior temporal areas (AC and pSTS). **b**, Prosodic representations occurred after pitch divergence (left grey rectangle) and extended from bilateral temporal areas to bilateral PMC and right IFG around and after stimulus offset (right grey rectangle).

**Table 1.** Cluster and peak statistics for phonemic and prosodic representations.

| Linguistic element | ROI | Cluster statistics |  | Peak statistics |  |  |
| --- | --- | --- | --- | --- | --- | --- |
|  |  | Extent (ms) | *p*-value | Latency (ms) | *t*<sub>28</sub> | Effect size (SEM) |
| Phoneme | L-AC | 72-228 | <0.001 | 156* | 7.456 | 0.685 (0.090) |
|  |  | 284-552 | <0.001 | 368 | 3.419 | 0.301 (0.087) |
|  |  | 564-644 | 0.0203 | 608 | 2.538 | 0.177 (0.069) |
|  | L-pSTS | 68-224 | <0.001 | 156* | 5.641 | 0.457 (0.080) |
|  |  | 312-404 | 0.0026 | 356 | 2.945 | 0.275 (0.091) |
|  | L-aSTS | 132-192 | 0.0018 | 160* | 3.773 | 0.284 (0.074) |
|  | R-AC | 68-212 | <0.001 | 108* | 6.667 | 0.595 (0.088) |
|  |  | 296-524 | <0.001 | 468 | 3.643 | 0.242 (0.065) |
|  |  | 532-592 | 0.0214 | 568 | 3.048 | 0.177 (0.057) |
|  | R-pSTS | 72-360 | <0.001 | 144* | 4.824 | 0.508 (0.103) |
|  |  | 392-500 | 0.0028 | 420 | 3.203 | 0.235 (0.072) |
|  | R-aSTS | 68-208 | <0.001 | 164* | 3.592 | 0.269 (0.074) |
| Prosody | L-AC | 280-580 | <0.001 | 416* | 4.147 | 0.351 (0.083) |
|  | L-pSTS | 332-532 | <0.001 | 412* | 3.662 | 0.297 (0.080) |
|  | L-aSTS | 264-316 | 0.0273 | 296 | 3.192 | 0.206 (0.064) |
|  |  | 432-556 | <0.001 | 456* | 3.754 | 0.324 (0.085) |
|  |  | 664-728 | 0.0132 | 712 | 3.015 | 0.211 (0.069) |
|  | L-PMC | 480-524 | 0.0297 | 488* | 2.669 | 0.159 (0.059) |
|  | R-AC | 300-644 | <0.001 | 408* | 3.845 | 0.466 (0.119) |
|  | R-pSTS | 316-560 | <0.001 | 496* | 4.638 | 0.428 (0.091) |
|  | R-aSTS | 380-564 | <0.001 | 408* | 3.855 | 0.326 (0.083) |
|  | R-PMC | 384-480 | 0.001 | 396* | 3.118 | 0.231 (0.073) |
|  |  | 528-592 | 0.004 | 560 | 3.186 | 0.207 (0.064) |
|  | R-IFG | 552-608 | 0.011 | 568* | 3.409 | 0.215 (0.062) |

This table lists statistics of significant clusters for phonemic and prosodic representations in the regions of interest (ROIs), along with their peak statistics. L-, left; R-, right; AC, auditory cortex; p/aSTS, posterior/anterior superior temporal sulcus; PMC, premotor cortex; IFG, inferior frontal gyrus; effect size, baseline-corrected Fisher-Z variance explained by both acoustic and categorical RDMS; SEM, standard error of the mean; \*, global peak of effect size in a given ROI.

Similar to phonemes, prosodic information was generally represented in bilateral ventral stream areas, but extended further into dorsal stream regions, including the bilateral PMC and right IFG (Fig. 3b). Prosodic representations emerged around 300 ms after stimulus onset, that is, after pitch contours started to clearly differ between the five prosody levels (184-250 ms, Supplementary Fig. 3; see also Supplementary Methods). The searchlight analysis confirmed both the extent and the timing of prosodic representations (Fig. 3b, Supplementary Fig. 2b). Comparable to phonemes, prosodic representations were strongly redundant in space and time, although there were subtle regional differences in when they emerged (first in the bilateral AC and pSTS, last in the right PMC/IFG and left pSTS) or peaked (first in the right PMC and right aSTS; Table 1). These subtle dynamics call for further analysis how prosodic information propagates within and between the ventral and dorsal streams.

### Transfer of phonemic and prosodic representations along ventral and dorsal streams

Next, we employed mTE analysis to elucidate how phonemic and prosodic information flows between the cortical areas identified above^36,37^. mTE is a multivariate extension of transfer entropy that allows the assessment of directed representational transfer between regions based on neural RDMs. This approach is similar to previous methods using Granger causality^40,41^ but offers advantages in capturing non-linear interactions.

Here, we focused on the subset of ROIs that showed either phonemic or prosodic representations. In these ROIs, we first re-generated time series of neural RDMs—this time without applying sliding windows—to ensure that the RDM at each time point was not influenced by preceding or subsequent time points. Specifically, we re-computed dissimilarities between MEG response patterns evoked by the five phoneme or prosody levels at each time sample from 0 to 800 ms after stimulus onset. Then, for each pair of ROIs, we quantified mTE between their time-resolved neural RDMs. We considered one ROI in a pair as the source and the other as the target at a time, and estimated mTE from the source to the target by lagging the target RDM series ranging from 4 to 300 ms. At each time lag, the measured mTE was normalised (mTEn) by the mean and standard deviation of the surrogate mTE distribution^42^, obtained by cyclically time-shifting the source RDM series^43^.

Fig. 4 shows representational transfer between ventral stream areas for phonemes (Fig. 4a-c, Supplementary Fig. 4a), and between both ventral and dorsal stream areas for prosody (Fig. 4d-f, Supplementary Fig. 4b). Propagation of phonemic information was extensive within and between hemispheres. Within each hemisphere, the AC and pSTS, that is, posterior temporal areas, were densely interconnected and projected anteriorly to the aSTS. The left (but not right) aSTS showed feedback connections to the ipsilateral AC and pSTS. Furthermore, the AC and pSTS generally projected to all contralateral areas, including their homologues. The only connection not identified was from the left AC to the right pSTS. The aSTS showed less interhemispheric connections, projecting only to its homologue.

**Fig. 4.**
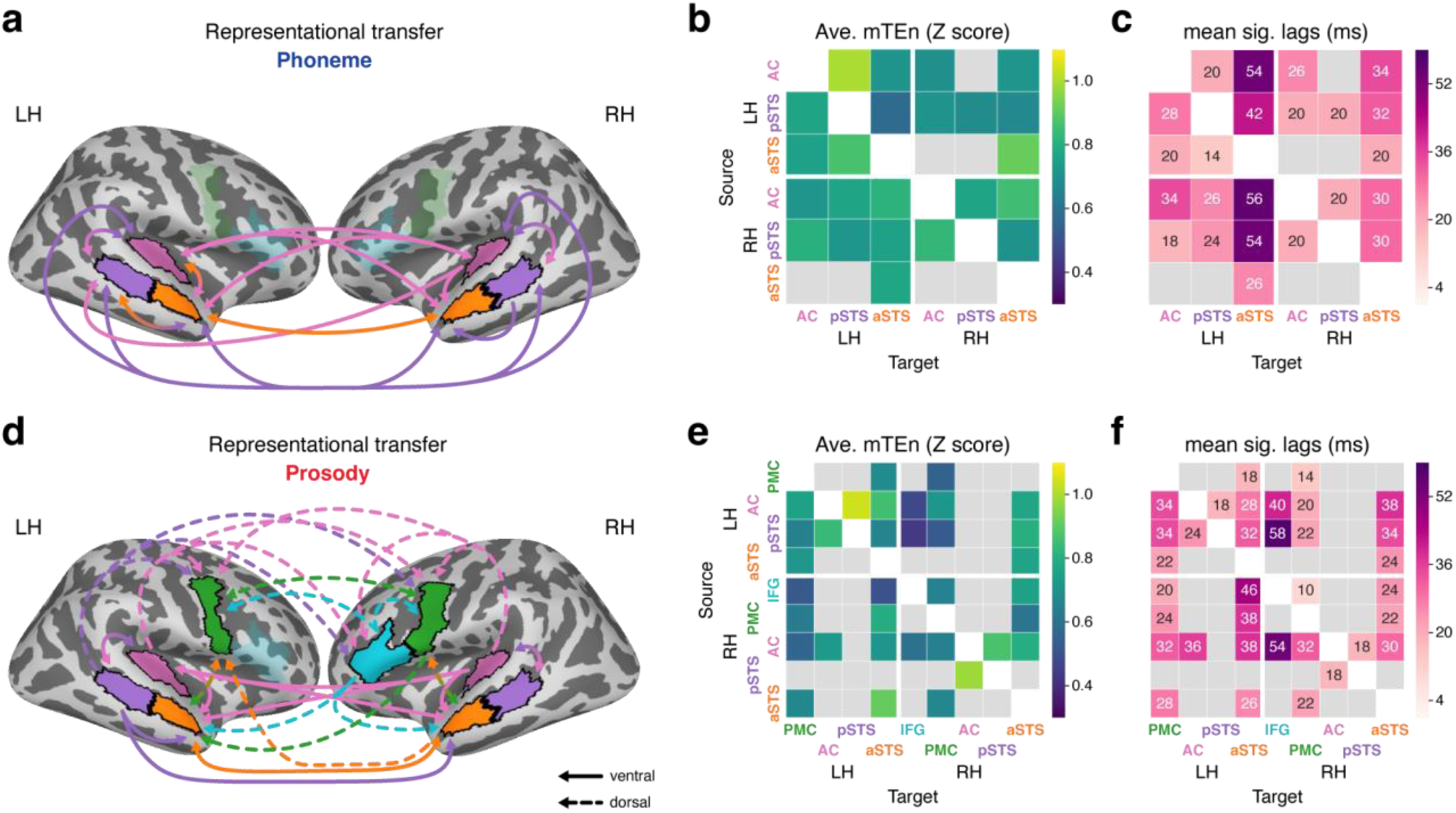
Transfer of phonemic and prosodic representations along ventral and dorsal streams. **a,d**, Directed representational transfer of phonemes (**a**) and prosody (**d**) between the ROIs that displayed significant corresponding representations (Fig. 3) in the left (LH) and right (RH) hemispheres (cluster-based permutation tests, one-sided *t*_28_-test on the normalised multivariate transfer entropy (mTEn) values against zero, CFT at *p* < 0.05, 100,000 permutations, FWER = 0.05 across time lags per directed connection, directed-connection-wise FDR *q* < 0.05). Arrows indicate directed representational transfer from the source to the target ROIs, colour-coded according to the source ROIs. **b,c,e,f**, Averaged mTEn (**b**,**e**) and mean time lags (**c**,**f**) across significant time lags, corresponding to **a** (**b**,**c**) and **d** (**e**,**f**). Rows and columns show the source and target ROIs, respectively (mTEn and significant intervals across time lags are shown in Supplementary Fig. 4). IFG, inferior frontal gyrus; PMC, premotor cortex; AC, auditory cortex; p/aSTS, posterior/anterior superior temporal sulcus. **a**, Phonemic representations were transferred along the ventral stream, from posterior to anterior temporal ROIs, with feedback connections from the left aSTS to the ipsilateral AC and pSTS. **d**, Prosodic representations propagated from posterior temporal ROIs to the aSTS along the ventral stream (solid) and to the bilateral PMC and right IFG along the dorsal stream (dashed). Note the relative dominance of the AC as a source compared to the pSTS (almost no projection from the right pSTS), and the cross-stream interaction between aSTS and PMC, two major target areas in the ventral and dorsal stream.

Interactions between posterior temporal areas tended to occur with shorter time lags (within about 50 ms), while their influence on the aSTS (especially, the left aSTS) lasted longer (up to 108 ms). These dynamics of phonemic representations are compatible with the established hierarchical organisation of the ventral stream with posterior-to-anterior information flow^12,14^.

Prosodic representations displayed partially similar ventral stream dynamics as phonemes, with some differences. Similar to phonemes, the AC and pSTS reciprocally exchanged prosodic information within each hemisphere, and the AC projected anteriorly to the aSTS in both hemispheres. However, pSTS-to-aSTS projections were confined to the left hemisphere, and no anterior-to-posterior feedback connections were found. Moreover, interhemispheric connections were relatively sparse: Exchange between posterior temporal areas was confined to one projection from the right AC to its left homologue, and only the bilateral AC (and the left pSTS) projected anteriorly to the contralateral aSTS. These dynamics of prosodic representations are, again, in line with posterior-to-anterior information transfer along the ventral stream^12,14^, but seem to highlight the relative importance of the AC (compared to the pSTS) as source.

In contrast to phonemes, prosodic representations were also transferred to dorsal stream areas. Especially posterior temporal areas, including the bilateral AC and left pSTS, projected dorsally to the bilateral PMC and right IFG, without recurrent connections. The left and right PMC communicated with each other, and both received top-down projections from the right IFG. Interestingly, the PMC and aSTS, major target areas in the dorsal and ventral streams, were bidirectionally connected within each hemisphere, and unidirectionally from right to left between hemispheres. The bilateral aSTS also received input from the right IFG. The transfer of prosodic representations took longer between more distant regions, with delays of up to about 90 ms, especially from posterior temporal areas in the ventral stream to the right IFG in the dorsal stream. Overall, these results demonstrate a strong cross-talk between the ventral and dorsal streams, with the aSTS and PMC serving as central hubs in the exchange of prosodic information.

### From acoustic to categorical representations of phonemes and prosody

After having shown how phonemic and prosodic representations are transferred along the ventral and dorsal streams, we examined whether and how these representations shift from acoustics to categories along these pathways and over time. To do so, we returned to the results of the time-varying RSA (Fig. 3) and partitioned the explained variance in each ROI, complemented by a whole-brain searchlight analysis. Given the high correlation between the acoustic and categorical RDMs (Supplementary Fig. 5, Supplementary Methods), we employed nested model comparisons to estimate the variance uniquely explained by each model RDM at a given time point, separately for phonemes and prosody (Fig. 2f).

Phonemes were represented acoustically in all ROIs along the ventral stream (Fig. 5a), while categorical representations were restricted to the bilateral AC and left pSTS (Fig. 5b). Both representations emerged early, within the first 200 ms after word onset, and overlapped in time. However, acoustic representations peaked consistently earlier (Table 2), suggesting a transition from acoustic to categorical representations over time^23,44^. These results were confirmed by the searchlight analysis, showing that acoustic representations extended more broadly and emerged slightly earlier in time than categorical representations (Supplementary Fig. 6a).

**Fig. 5.**
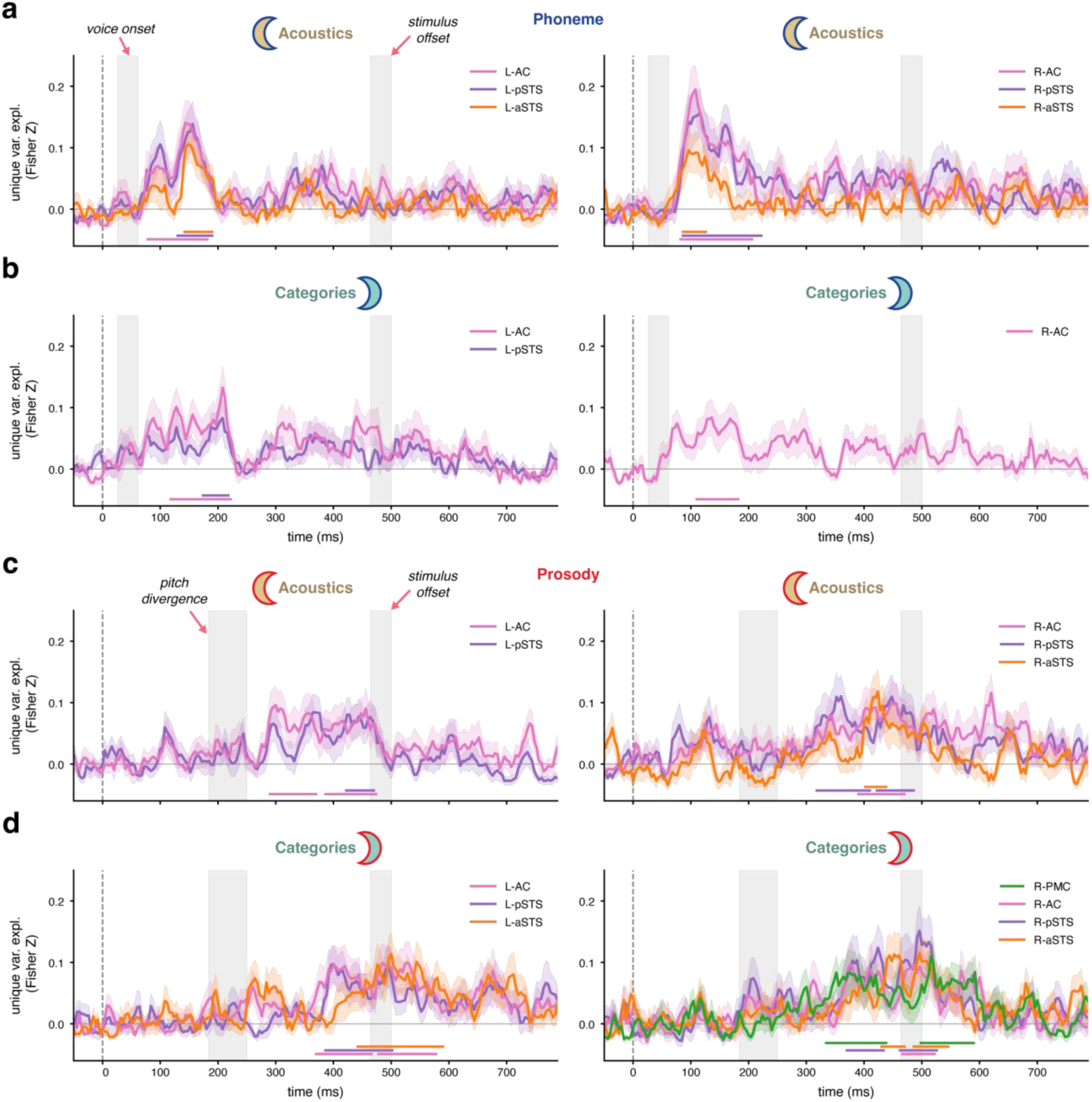
Acoustic and categorical representations for phonemes and prosody in ventral and dorsal stream regions. **a-d**, Time-varying variance (Fisher-Z scale, baseline-corrected) uniquely explained by the acoustic (**a**,**c**) or categorical RDM (**b**,**d**) in the left-(L-; left panel) and right-hemispheric (R-; right panel) ROIs that exhibit significant unique representations of phonemes (**a**,**b**) or prosody (**c**,**d**). Statistical significance was determined by cluster-based permutation tests (one-sided *t*_28_-test against zero, CFT at *p* < 0.05, 100,000 permutations, FWER = 0.05 across time points per ROI, ROI-wise FDR *q* < 0.05). Shaded areas indicate SEM. Horizontal lines above the x-axis show significant time intervals (ROI-wise cluster and peak statistics are summarised in Table 2). PMC, premotor cortex; AC, auditory cortex; p/aSTS, posterior/anterior superior temporal sulcus. **a**, Acoustic representations of phonemes emerged after voice onset (left grey rectangle) and extended across bilateral temporal ROIs. **b**, Phoneme categories were represented slightly later, but overlapped considerably in time with the acoustic representations (**a**). Notably, the categorical representations were distributed less broadly than acoustics, limited to the bilateral AC and left pSTS. **c**, Prosody acoustics were represented after pitch divergence (left grey rectangle) in all bilateral temporal ROIs except for the left aSTS. **d**, Successive categorical representations of prosody overlapped with the acoustic representations (**c**) in time but lasted longer beyond stimulus offset (right grey rectangle). Note that categorical representations of prosody were spatially broader than those of phonemes (**b**), including the bilateral aSTS and right PMC.

**Table 2.**
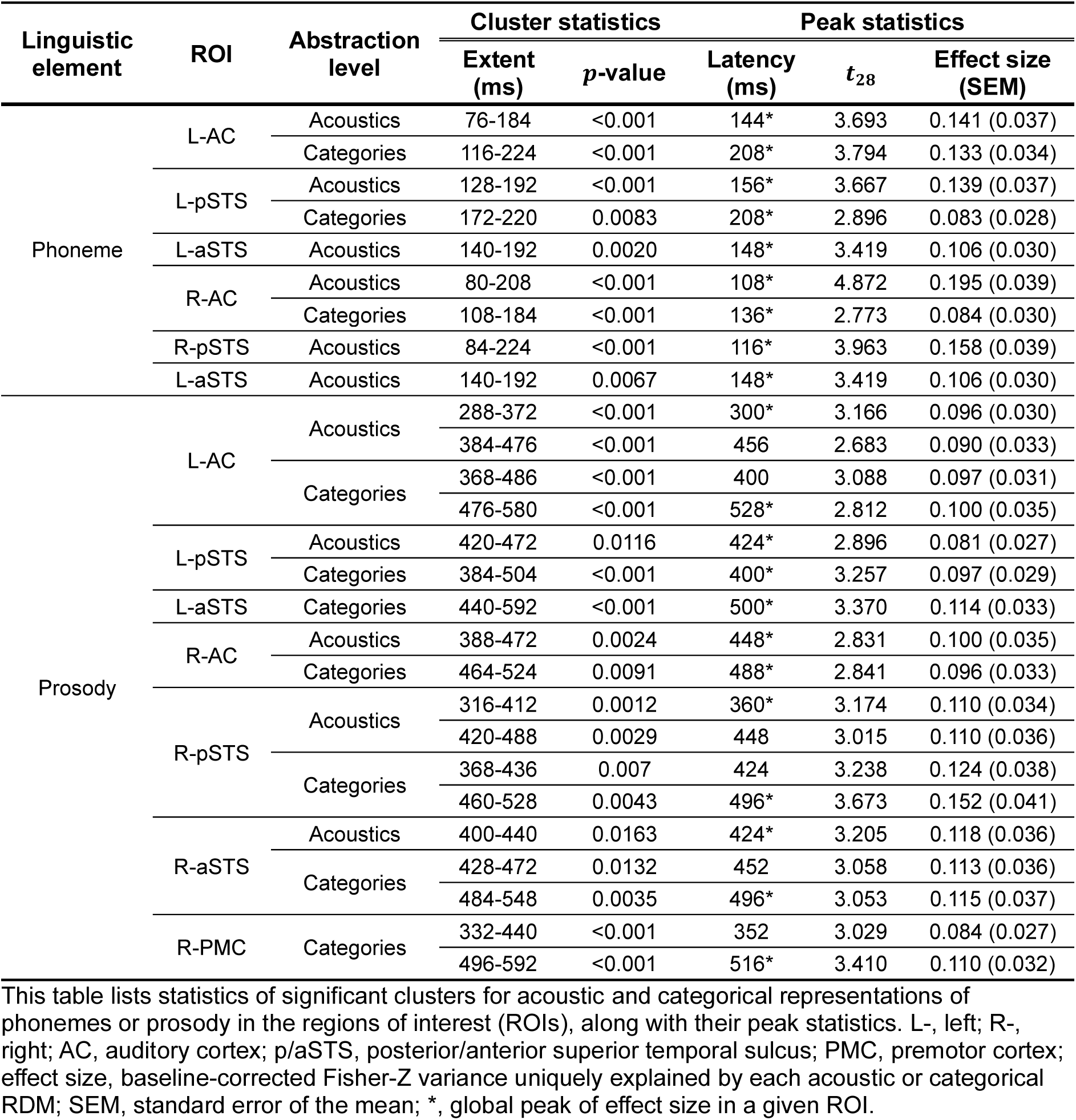
Cluster and peak statistics for acoustic and categorical representations of phonemes and prosody.

Prosody, similar to phonemes, was represented acoustically in all temporal ROIs, except for the left aSTS (Fig. 5c). However, categorical representations were spatially more widespread for prosody than phonemes, including all ventral stream areas as well as the right PMC (Fig. 5d). Notably, categorical representations also peaked nominally later (Table 2), similar to phonemes, and lasted longer than acoustic representations, in line with a gradual transfer from acoustics to categories^23,44^, although acoustic and categorical representations showed considerable overlap in time. Interestingly, the right PMC exhibited only categorical, not acoustic, representations of prosody, in two time windows: one before and the other after stimulus offset. Again, all results were confirmed by the searchlight analysis (Supplementary Fig. 6b).

Additionally, we compared the time-varying variance uniquely explained by acoustic or categorical RDMs between homologous ROIs to explore the lateralisation of phonemic and prosodic representations (Supplementary Fig. 7,8). None of the representations was significantly lateralised, neither for phonemes nor prosody (all cluster *ps* > 0.05).

Altogether, both the phoneme and prosody data suggest a fast transformation of acoustics into categories in the temporal lobe^23,44^. However, both abstraction levels were sustained in parallel over time, in line with recent evidence showing preserved lower-level representations during higher-level processing stages^23^. Meanwhile, the spatial dynamics of this transformation in the ventral and dorsal streams differed between phonemes and prosody, suggesting partially different mechanisms of abstraction.

### Task modulation effect on phonemic and prosodic representations

Lastly, we tested whether task demands to categorise phonemes or prosody intensified the respective neural representations. This idea builds on earlier studies showing that task demands can modulate cortical activity and enhance the encoding of task-relevant information, even when the sensory input remains unchanged^45,46^. To this end, we split the data by task (Fig. 1b), and estimated and compared the time-varying variance explained by the model RDMs in each ROI that had represented either phonemic or prosodic information (Fig. 3).

While the prosody task did not enhance any prosodic representations (all cluster *ps* > 0.05), the phoneme task enhanced the phoneme representations, selectively in the left pSTS, compared to the prosody task, around voice onset, peaking at 52 ms post-onset (Fig. 6a; *t*_28_ = 3.334, baseline-corrected Fisher Z variance contrast = 0.228, standard error of the mean (SEM) = 0.067). No such effect was found in other ROIs (all cluster *ps* > 0.05).

**Fig. 6.**
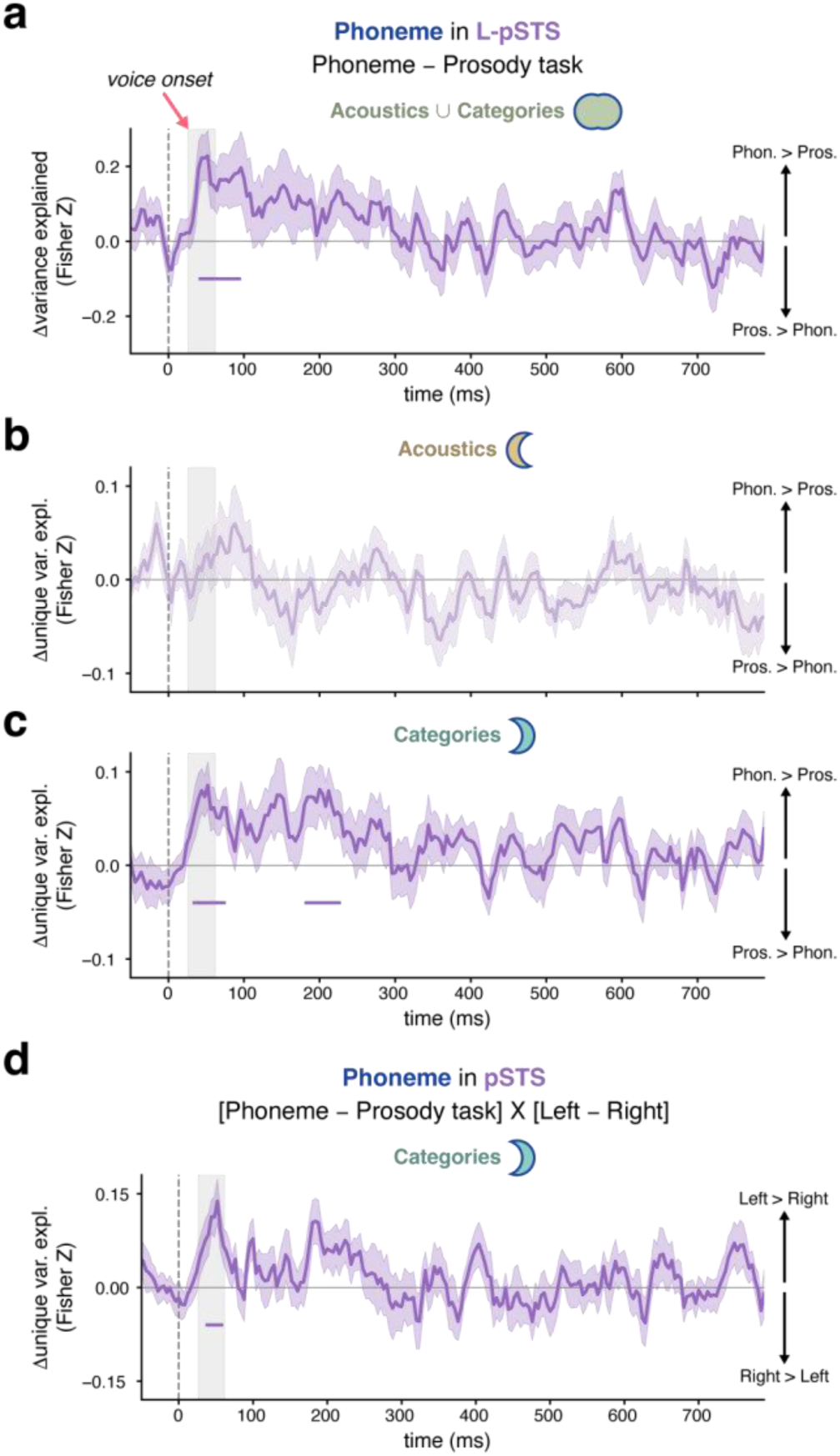
Task influences on phonemic representations in the left posterior superior temporal sulcus (pSTS). **a-c**, Contrasts of time-varying variance (Fisher-Z, baseline-corrected) explained by both acoustic and categorical RDMs for phonemes (**a**) or uniquely explained by either RDM (**b**,**c**) between the phoneme and prosody identification tasks in the left pSTS. Shaded areas indicate SEM. Horizontal lines show significant time intervals (cluster-based permutation tests, one-sided *t*_28_-test, CFT at *p* < 0.05, 100,000 permutations, FWER = 0.05 across time points). **a**, Phonemic representations were enhanced during the phoneme tasks (correction for multiple ROI testing at FDR *q* < 0.05) around voice onset (grey rectangle). **b,c**, The phoneme task intensified the categorical representations (**c**), but not acoustic representations (**b**, pale). **d**, Contrast of task-modulated unique variance for the categorical RDM (Fisher-Z) between the left and right pSTS over time (statistical inference and plotting details as above, but with two-sided *t*_28_-test). Categorical representations of phonemes were selectively enhanced in the left pSTS.

Next, we investigated which abstraction levels of phonemic representations in the left pSTS were modulated by task demands. Using variance partitioning, we compared the acoustic or categorical representations of phonemes in the left pSTS between the two tasks. While the acoustic representations were not modulated by task (Fig. 6b), the categorical phoneme representations were amplified during the phoneme task (Fig. 6c). This effect was evident again around voice onset (peak at 52 ms, *t*_28_ = 3.345, baseline-corrected Fisher Z unique variance contrast = 0.086, SEM = 0.025) and around 200 ms post-onset (peak at 200 ms, *t*_28_ = 3.024, baseline-corrected Fisher Z unique variance contrast = 0.081, SEM = 0.026).

Finally, we compared the task-modulation effect between the left and right pSTS. As depicted in Fig. 6d, the categorical representations of phonemes were enhanced by the phoneme task significantly more strongly in the left than right pSTS, again around voice onset, peaking at 52 ms (*t*_28_ = 3.914, baseline-corrected Fisher Z unique variance contrast = 0.138, SEM = 0.035). Overall, these findings highlight the special role of the left pSTS in categorical phoneme perception and its modulation by task focus, in line with the literature^13,22,46^.

## Discussion

Successful speech perception requires transforming continuously variable acoustic signals into perceptually distinct categories. Here, we studied the neural basis of this perceptual transformation for phonemes and prosody, two fundamental building blocks of speech that rely on different acoustic cues and contribute to different aspects of linguistic meanings.

Using behavioural psychophysics and MEG, combined with time-resolved RSA and mTE analysis, we found that both phonemic and prosodic representations underwent an acoustic-to-categorical abstraction along the ventral stream, with additional involvement of the dorsal stream for prosody. However, contrary to predictions of classical hierarchical processing models, the two abstraction levels significantly overlapped in time and space, indicating their parallel rather than serial processing. The convergence of this finding for both linguistic elements points to a shared principle of representational abstraction. On the other hand, while acoustic representations of phonemes and prosody were similarly distributed across the temporal cortex, their categorical representations markedly differed in spatial extent: Phoneme categories were represented in bilateral posterior temporal areas, with enhanced representations in the left pSTS under top-down focus on phonemes. In contrast, prosody categories were more widespread, including the bilateral aSTS and right PMC. These results suggest partially distinct abstraction mechanisms within the dual streams depending on the type of cue and linguistic function of phonemes and prosody.

To start with, our findings provide empirical evidence that prosody, like phonemes, is perceived categorically. While the categorical perception of phonemes is well established at both behavioural and neural levels^15,17,20,29,33^, clear support for prosody has been lacking.

Several studies have suggested that prosodic pitch accents are neurally encoded as discrete categories^21,26^. However, these findings did not explicitly dissociate categories from stimulus acoustics and were not directly linked to listeners’ perception (i.e., behaviour), leaving it unclear whether this encoding genuinely reflects the categorisation process. Here, we demonstrate the categorical perception of question-statement prosody using behavioural psychophysics (Fig. 1d,e), and identify the neural signatures of their perceived categories beyond the relevant acoustic cues (Fig. 5d). These results are a first step towards substantiating the psychological reality of prosody categories, beyond phonemes, as fundamental building blocks of speech.

The representational dynamics of phonemes and prosody we observed challenge classical hierarchical models of speech perception which posit a serial encoding—first of acoustic cues in sensory areas, and then of abstract linguistic categories in higher-order regions^6,9–11^. While we did observe an acoustic-to-categorical abstraction in time (Fig. 5) and a posterior-to-anterior propagation (Fig. 4) of phonemic and prosodic representations along the ventral and (dorsal) streams, our findings deviated from serial hierarchical processing models in three respects.

First, we found that perceived categories of both phonemes and prosody were already encoded in the AC (Fig. 5b,d). This suggests that abstract linguistic representations may emerge at earlier stages of cortical auditory processing than predicted by the traditional models^19–21^. Note that it is unlikely that these categorical representations in the AC merely reflect top-down perceptual biases from higher-order areas^47,48^. Even though we did observe representational transfer from the pSTS to the AC for both linguistic elements, and from the aSTS to the AC in the left hemisphere for phonemes (Fig. 4), the categorical representations mostly appeared earlier in the AC than in these other regions (Fig. 5b,d). Rather, our data support the proposal that early sensory areas are more than mere auditory feature detectors, but are capable of representing auditory objects^49,50^. For example, a recent ultra-high-field neuroimaging study has shown that responses in superficial layers of primary auditory areas go beyond simple auditory analysis in deeper layers, and are best explained by a processing complexity comparable to that of non-primary auditory areas^51^. Whether such processing capacity underlies the categorical representations of abstract linguistic elements observed in the AC can be tested in future studies with laminar resolution.

Second, we found spatially distributed acoustic representations of both phonemes and prosody across the ventral stream (Fig. 5a,c). These results are inconsistent with predictions of serial hierarchical processing models, which assume auditory analysis is confined to early sensory areas^6,10,11^. Similar findings from intracranial recordings show that acoustic-phonetic features are redundantly encoded throughout the temporal cortex alongside higher-level linguistic information^23^. This suggests that acoustic details are not discarded after abstraction but are retained in higher-order regions, potentially supporting robust speech perception by enabling the retrieval of earlier acoustic information for re-analysis^23,52^.

Lastly, our findings revealed that phoneme and prosody categories were represented largely (although not fully) in parallel with their acoustic cues, both in time and space (Fig. 5). This challenges serial hierarchical processing models, which predict that speech categorisation occurs after auditory processing and primarily in higher-order regions^6,9,11^.

Both categorical and acoustic representations emerged shortly after relevant acoustic events (∼200 ms), and although categories occurred nominally later, both representations were sustained in parallel over time. Moreover, categorical representations spatially overlapped with acoustic representations across temporal areas, especially in the AC and pSTS for both prosody and phonemes. This seems to suggest a special role of the posterior temporal cortex in the acoustic-to-categorical transformation, in line with prior studies that have indicated this rather focal region as site of the actual categorisation operation, in particular for phonemes^29^, but also prosodic pitch accents^21^, and linked to categorical decisions^33^.

Interestingly, the processing of phonemes and prosody displayed subtle differences in the AC and pSTS. While prosodic representations were unaffected by task demands, top-down efforts to identify phonemes selectively enhanced their categorical representations in the left pSTS (Fig. 6). This indicates the relative importance of the left pSTS in phonemic processing^13^, consistent with prior evidence that the left pSTS primarily encodes phonetic features and categories of phonemes, and plays a crucial role in spoken word recognition^22,24,29^. In contrast, the AC seemed to play a more central role than the pSTS for prosody. The AC is known to be sensitive to pitch-related features^21,22^, which may underlie its more dominant role in sending prosodic information (Fig. 4). These findings may suggest differential weightings of the AC and pSTS to support the specific processing demands of different linguistic elements, in line with recent observations from direct cortical recordings^21,22^.

On top of these subtle differences in the AC and pSTS, prosody categories extended markedly more anteriorly along the ventral stream than phoneme categories, including the aSTS (Fig. 5b,d). This difference may stem from general tuning preferences of neuronal populations in the temporal cortex, which process auditory information at different timescales, with gradually increasing temporal integration windows along the posterior-anterior axis of the temporal lobe^53,54^. This cortical gradient is accompanied by a shift in neuronal encoding properties—from spectrotemporal cues to semantic categories of sounds^53^. Overall, both properties align with characteristic differences between phonemes— which are brief and often lack concrete meanings^2,32^—and prosody—which generally spans longer durations and imparts specific meanings such as the speaker’s intention^3^. This suggests that general neurofunctional gradients may be differently engaged by different linguistic elements, in a way that prosody categorisation reaches further along these cortical gradients.

Our findings further illuminate the role of the dorsal stream in speech processing. A dominant framework proposes that motor and inferior frontal regions support speech perception by mapping auditory information onto articulatory-motor representations^55,56^ or by contributing to decision-making^17,18^. However, it is often argued that these regions are generally not involved in sub-lexical speech processing but are instead recruited under specific demands, such as task engagement or noisy listening conditions^6^. Our data suggest that the dorsal contribution differs depending on the type of linguistic elements tested. While we found no phonemic representations along the dorsal stream (Fig. 3a), prosodic representations extended to the bilateral PMC and right IFG (Fig. 3b). More specifically, the right PMC represented prosody categories during two time windows, before and after stimulus offset (Fig. 5d), implying a motor contribution to prosody categorisation^7^ across multiple processing stages^56^. The right IFG displayed unspecific acoustic-categorical representations after word offset (Fig. 3b, Fig 5c,d), potentially involved in decision-making based on shared acoustic and categorical information^11^. Importantly, this disparity between phonemes and prosody is unlikely to result from differences in perceptual or processing demands^6^, as both involved the same identification task and were matched in difficulty. One possible explanation is that dorsal involvement is stronger for linguistic elements unfolding over longer timescales. Supporting this idea, previous MEG evidence shows that auditory-motor interaction is restricted to rates up to about 5 Hz (>200 ms)^57^, which may provide a suitable temporal window for processing prosody but could be too slow to convey sub-lexical information such as VOT. This temporal constraint may underlie the observed engagement of the dorsal stream in prosodic, but not phonemic, processing.

What is more, prosodic processing involved extensive representational transfer between the ventral and dorsal streams (Fig. 4d-f). More specifically, the PMC received projections from posterior temporal areas and was bidirectionally connected to the aSTS. These links may underlie auditory-motor integration supporting the categorical perception of prosody^55,56^. Similarly, the right IFG received afferent input from posterior temporal areas and back-projected to the bilateral PMC and aSTS, possibly modulating their representations^58^.

Together, these findings suggest that the PMC and aSTS serve as central hubs for cross-stream interactions in the exchange of prosodic information. Given that there is no direct anatomical connection between the PMC and aSTS^7,8^, future studies should examine whether the observed functional links are indirectly driven by common input from posterior temporal areas and the right IFG^59^, or possibly mediated by subcortical regions^60^.

Apart from the ventral-dorsal cross-talk for prosody, both phonemic and prosodic representations were characterised by extensive interhemispheric transfer (Fig. 4). This may account for the absence of clear lateralisation effects (Supplementary Fig. 7,8), despite previously reported hemispheric processing asymmetries^7,8,13^.

Before closing, several caveats regarding our findings should be noted. First, we examined only a single example of a phonemic and a prosodic cue—/b/-/p/ along the VOT continuum, and statement-question pitch contours—limiting the generalisabilty of our findings to other distinctive features. However, these exemplars, that deliberately tap into different timescales to probe and extend prevailing knowledge, already revealed aspects of speech processing that cannot be accounted for by existing models, such as categorical representations in early sensory areas and parallel acoustic and categorical processing in both time and space. Future studies on other phonemic and prosodic cues will help to substantiate the observed representational dynamics and further estimate the extent to which traditional serial hierarchical processing models need to be revised. Second, our mTE analysis assessed the overall information flow of phonemic and prosodic representations, leaving unresolved which levels of abstraction are transferred between regions over time.

Additionally, some of the observed connections might have arisen from common sources or reflected secondary links via intermediate areas^59^. Future studies should address these potential indirect connections to clarify the precise transfer of phonemic and prosodic representations, with taking different levels of abstraction into account. Lastly, we investigated phonemes and prosody in isolation, offering limited insight into how different linguistic elements integrate—a process crucial for speech comprehension. Future work should explore the interaction between phonemic and prosodic information in the brain to better understand how a unified representation of speech is formed.

In sum, we demonstrate the dynamic propagation and acoustic-to-categorical transformation of phonemes and prosody along the ventral and dorsal speech streams. However, our results do not fully conform with the “serial” view of classical hierarchical processing models^6,9–11^. Instead, they point to a shared principle of parallel acoustic and categorical processing over time and across space for both linguistic elements, potentially promoting robust speech perception. The partially different dynamics and distributions of phonemes and prosody along the ventral and dorsal streams indicate somewhat distinct mechanisms of abstraction, arising from the physical properties and linguistic functions of the respective elements. This flexible and complementary adaptation of regional weights and information flow to transient phonemic and sustained pitch-based prosodic cues is likely to underlie listeners’ ability to concurrently process and grasp the richness of meanings conveyed by different linguistic elements in the continuous speech stream.

## Methods

### Participants

34 native German speakers (17 females, mean age = 26, SD = 4) were recruited and provided written informed consent prior to participation. All participants self-reported normal hearing with no history of neurological or psychiatric disorders, and were assessed as right-handed by the Edinburgh Handedness Inventory (median laterality quotient = 95, 10^th^-90^th^ percentiles: 80-100)^61^. Five participants (two females) were excluded due to poor behavioural performance or low MEG signal quality, leaving 29 participants for the analyses. This study adhered to the ethical standards of the Declaration of Helsinki and was approved by the Ethics Committee of the Medical Faculty, Leipzig University (403/14-ff).

### Stimuli

Stimuli consisted of the German words “Bar” and “Paar,” which gradually varied in both phonemic and prosodic cues. Mono-syllabic words were used to minimise the involvement of compositional linguistic processing, such as syntactic and semantic processes^7^. For stimulus development, we first recorded “Bar” and “Paar” with both statement and question intonations, produced by two German native speakers (one female) in a soundproof booth using a Røde NT55 microphone and Audacity® (version 2.0.5, sampling at 16 kHz, 16-bit, mono). We then segmented the recordings into word-initial phonemes (/b/ and /p/) and word stems, and used them to separately morph the VOT and pitch contour for each speaker in MATLAB (R2014a, MathWorks).

We first morphed the pitch contour of the word stems from statement to question using STRAIGHT^62^. For this, we placed anchor points on corresponding time-frequency landmarks in the two utterances (see Sammler et al.^7^), and decomposed the sound signals into five features (*F*_0_, frequency structure, duration, spectrotemporal density and aperiodicity). We then re-synthesised new word stems by interpolating these features in 2% steps on linear (duration and aperiodicity) and logarithmic (*F*_0_, frequency structure and spectrotemporal density) scales. The resynthesis continued up to 120% of the original question by extrapolating the landmarks, resulting in 61 word stems with a duration of 430 ms. Next, we morphed the VOT by manually splicing incrementally longer segments (1-ms steps) of aspiration noise from the original /p/ between the plosive (/b/) and the word stems in 61 steps, starting from the original /b/ (0-ms VOT) reaching up to 60-ms VOT. The splicing produced 61 VOT morphs, applied to each of the 61 word stems. This resulted in 3,721 stimuli for each speaker (7,442 stimuli in total), varying along 61 x 61-step orthogonal morphing continua of VOT and pitch contour (middle panel of Fig. 1). Additionally, we inserted a 10-ms silence at the beginning of the stimuli and applied a 10-ms fadeout at the end to prevent unwanted clicks. All stimuli were normalized in root-mean-square amplitude.

Out of the 61 x 61 steps, 5 x 5 morph levels were selected for each participant based on two behavioural pre-tests (see Supplementary Methods), leading to a subset of 25 stimuli per speaker (50 stimuli in total) presented during the MEG experiment. The first pre-test used an adaptive-staircase method to identify five levels of VOT and pitch contour centred on individual category boundaries. The distance between the five levels was selected to ensure that the difficulty of categorising phonemes and prosody was matched. Stimulus selection was validated in the second pre-test and adjusted, if necessary. This approach ensured that stimuli covered the feature-range that gave rise to the acoustic-to-categorical transition in phonemes and prosody in each speaker and participant, irrespective of the absolute acoustic values.

### MEG experiment procedure

During the MEG experiment, stimuli were delivered via air-conduction earphones (ER3-14A/B, Etymotic Research) at an average of 74 dB SPL. The volume was adjusted individually based on each participant’s hearing threshold, measured before the experiment proper using a stimulus that was not presented later.

The experiment consisted of six runs, each of which contained four blocks. Across the four blocks, participants were alternately asked to identify either the word (“Bar”/“Paar”) or the prosody (“statement”/“question”) of the stimuli (Fig. 1b). The stimuli of the same speaker were presented in two blocks in a row per run, but with different tasks. For example, all stimuli of the female speaker were first presented with a phoneme, then with a prosody task, followed by a phoneme and a prosody block with male stimuli. The order of tasks and speakers per run was randomised and counterbalanced across participants.

In each block, participants listened to stimuli in separate trials. Each block began with four practice stimuli featuring extreme levels of phonemes and prosody. These practice trials were intended to help participants adapt swiftly to the task and were excluded from the analyses. After the practice trials, 25 unique stimuli (i.e., 5 x 5 morph levels of the speaker) were presented pseudorandomly following the principles of a type-1 index-1 sequence to prevent carry-over and position effects^63^. Specifically, the stimuli were ordered such that, ideally, each morph level was followed by every other morph level, including itself, twice per block. Stimulus order varied across speakers and runs.

During stimulus presentation, participants were instructed to carefully listen to each word and mentally categorise whether the speaker said “Bar” or “Paar” or uttered “statement” or “question,” depending on the task. The task was indicated on screen at the beginning of each block and the colour of the fixation cross during the block reminded participants of the type of task (green for phoneme and red for prosody). Participants were asked to report their percept by pressing a button on a response box corresponding to the on-screen category labels, but only in one-sixth of the trials (i.e., 10 responses for each of the five phoneme and prosody levels per speaker). This was to maximise the overall number of trials (and signal-to-noise ratio) within the recording session, by saving time spent on responses and pause intervals, and while acquiring sufficient behavioural data for fitting reliable psychometric functions. Response trials occurred randomly within each block to ensure participants remained attentive throughout the experiment. Additionally, response buttons were randomly assigned on each response trial to avoid confounding category labels with preparatory motor activity. The labels “Bar” and “Paar” (or “statement” and “question”) were presented on either the left or right side of the screen after stimulus offset, with their positions balanced across stimuli, tasks, and speakers.

Stimulus onset asynchrony was 2800 ms with a random jitter between −500 and 500 ms. Response trials included an additional 2000-ms timeout for the button press, and button presses were followed by a 1000-ms pause before the onset of the next stimulus. The entire procedure was controlled by Presentation® software (version 18.0, Neurobehavioral Systems) and lasted about two hours, including optional breaks between the runs.

### Behavioural analysis

For each participant, we computed the proportions of “Paar” and “question” responses at the five phoneme and prosody levels for each speaker across the six runs. Four participants in whom the difference between minimal and maximal proportion was smaller than 0.6 in at least one of the tasks were excluded from the analysis, as they either perceived no clear question/Paar or no clear statement/Bar.

The proportions at the five morph levels were then used to fit psychometric curves to each participant’s data per task and speaker using the following expression:

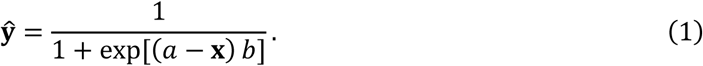

***x*** and ***ŷ*** are column vectors, representing the five levels and the estimated proportions, respectively. *α* is the point where the curve predicts equal proportions for both categories, and *b* is the slope of the curve at *α*. The fitted 50%-point *α* and slope *b* were taken as the individual estimates of PSE and perceptual difficulty. The PSE and perceptual difficulty were separately compared between tasks and speakers at the group level using repeated measures analyses of variance (rmANOVAs). As no speaker-related effects were found for both measures, we averaged the behavioural responses across speakers for the subsequent analyses.

A further analysis was conducted to verify the categorical perception of phonemes and prosody. For each task, we fitted a linear function ***ŷ*** = *m**x*** + *n*, and a psychometric curve, as in Equation (1), to each participant’s responses, and measured the adjusted *R*^2^ as a metric for their goodness of fit. The adjusted *R*^2^ between the linear and curve fits was compared at the group level using an rmANOVA, with function (linear vs. curve) and task (phoneme vs. prosody) as factors, to assess whether the increase of response proportions across the five levels better followed a psychometric curve rather than a linear function for both phonemes and prosody.

### MEG data acquisition and preprocessing

MEG data were collected during each run using a Neuromag Vectorview system (Elekta) with 306 sensors (102 magnetometers and 204 gradiometers). The MEG signals were sampled at a rate of 1,000 Hz, with a passband from DC to 330 Hz, while participants’ head positions were tracked using five head coils.

The acquired MEG data were preprocessed using MNE-Python (version 1.1.0). For each run, external interference was suppressed by spatiotemporal signal-space separation, including correction for head movements and semi-automatically identified noisy channels (median = 3 sensors, 10^th^-90^th^ percentiles: 2-5). The signals were then lowpass-filtered at 80 Hz using a finite impulse response (FIR) filter with a 10-Hz transition bandwidth (one-pass, zero-phase shift, non-causal, Hann window), followed by the reduction of line noise (50 Hz) and its harmonics using ZapLine^64^. The low-frequency range below 0.3 Hz was further attenuated with a highpass filter (a one-pass, zero-phase-shift, non-causal FIR filter using a Hann window with a transition bandwidth of 0.3 Hz) that was designed to replace the need for baseline correction^65^. The filtered signals were downsampled to 250 Hz and subjected to independent component analysis (ICA) to remove artefacts related to physiological noise, such as eye movements and heartbeats. The ICA solution was derived from the data concatenated across all six runs for each participant. After the artefact removal, the signals were segmented into epochs from −200 to 800 ms around stimulus onset. Epochs with excessive noise were discarded by Autoreject^66^, after which one participant was excluded for having more than 20% of epochs rejected. On average, 1,191 epochs (SD = 44) were retained for each participant across the six runs.

### Source reconstructions

Source activity was reconstructed from the preprocessed MEG signals with anatomical constraints, derived from high-resolution individual T1-weighted images (1 x 1 x 1-mm^3^ voxel size) obtained using a 3-T magnetic resonance imaging (MRI) scanner (Siemens).

Anatomical images were segmented using FreeSurfer (version 7.3.2) to construct a volume conduction model and source space for each participant. The volume conduction was estimated by a single-layer boundary element model created with the watershed algorithm applied to the inner skull. The source space comprised 10,242 vertices per hemisphere along the cortical surface, where dipole orientations were defined as the surface normal with a loose constraint. The anatomical image was aligned with the sensor space by coregistering the individual MEG and MRI data through iterative procedures based on the fiducials and digitised points on the head model. The sensor signals were then projected onto the source space using eLORETA^67^ (regularisation parameter = 1/9), while being whitened by a covariance matrix of sensor noise. The noise covariance matrix was estimated from the prestimulus period (−200 to 0 ms around stimulus onset) of all epochs and regularised by the Ledoit-Wolf method with cross-validation^68^. Source reconstructions were performed on the evoked responses to each of the five phoneme or prosody levels, averaged across the corresponding epochs for each speaker and run when estimating task-agnostic source activity, as well as for each task when estimating task-specific source activity. The estimated source activity in the individual source space was finally morphed into the common *fsaverage* space for subsequent analyses.

### Time-resolved RSA

Time-resolved RSA was employed to identify where and when phonemes and prosody were represented in the brain and at which level of abstraction, that is, in terms of their acoustics or categories^34^. This analysis involved constructing RDMs based on multivariate neural dissimilarities in specific regions and across time, reflecting the relationship between stimuli in the local representational space. To delineate the characteristics of the representational structure, neural RDMs were compared with model RDMs built on acoustic and perceptual dissimilarities of the stimuli, respectively. We performed time-resolved RSA in individual participants and analysed phoneme and prosody separately.

Neural RDMs were primarily constructed from source activity in 10 predefined ROIs (Fig. 2a). The ROIs were defined based on a multimodal cortical parcellation^69^ to be comparable in size and spatially distinct to minimise cross-talk. They covered areas previously shown to be involved in the processing of phonemes and prosody^7,8,28^, including the bilateral IFG (44, 45, IFSa, IFSp), PMC (6r, 6v, PEF, 55b), AC (A1, Mbelt, LBelt, PBelt), pSTS (STSdp, STSvp) and aSTS (STSda, STSva). The ROI-based approach was complemented by a whole-brain searchlight analysis (Fig. 2a), in which a searchlight with 10-mm radius was shifted across all vertices in the source space. For both approaches, sliding time windows of 24 ms were used to generate time series of neural RDMs in specific regions. Time windows advanced in 4-ms steps for the ROIs, while half-window-sized steps were applied in the searchlight analysis to reduce computational load. In each ROI or searchlight location, we extracted task-agnostic MEG source activity patterns evoked by the five phoneme or prosody levels within a given time window for each speaker and run, and measured the squared Euclidean distance between all level pairs within each speaker, cross-validated across the six runs^39^ (Fig. 2b). The measured neural dissimilarities were then averaged across speakers, yielding a time series of 5 x 5 neural RDMs per region and participant, separately for phonemes and prosody (Fig. 2c).

Meanwhile, model RDMs were built for stimulus acoustics and perceived categories (Fig. 2d). Stimulus acoustics were coded as the original morph steps of the stimuli along the 61-step phoneme or prosody continuum for each speaker, which preserved the statistics of stimulus acoustic features (see Supplementary Fig. 9, Supplementary Methods). Perceived categories were modelled as the expected proportion of “Paar” and “question” responses at each of the five morph levels, derived from the psychometric curves fitted to individual behaviour pooled across speakers. Both acoustic and categorical dissimilarities between the five phoneme or prosody levels were estimated by calculating the squared Euclidean distance between the respective values, producing 5 x 5 RDMs, each normalised to a range between 0 and 1. Finally, we averaged two RDMs encoding acoustic dissimilarities from the different speakers, resulting in one acoustic and one categorical RDM per participant for both phonemes and prosody (Fig. 2e).

We quantified neural representations of phonemes and prosody by nested model comparisons. Specifically, three types of nested linear models were built to predict each time step of the neural RDMs in a given ROI or searchlight, as follows:

- Full model: neural RDM ∼ intercept + acoustic RDM + categorical RDM
- Reduced model: neural RDM ∼ intercept + acoustic OR categorical RDM
- Null model: neural RDM ∼ intercept

Before entering the linear models, all RDMs were vectorised based on the lower triangular entries, as they were symmetric. The model fits (*R*^2^) were estimated by fitting Equation (2) below using non-negative least squares (NNLS), which prevented negative weights that could otherwise arise from negative distances in the neural RDM due to cross-validation:

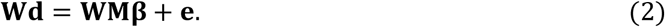

***W*** is a pre-whitening matrix multiplied to both the neural RDM column vector ***d***, and the predictor matrix ***M***. ***β*** is a column vector of weights to be estimated, corresponding to the number of predictors. ***e*** is the error term. The pre-whitening matrix ***W*** was introduced to mitigate the problem that the distance estimates in the RDMs were intrinsically correlated due to the shared morph levels among them, leading to sub-optimal comparisons between the neural and model RDMs^39^. Thus, we constructed ***W*** from a noise covariance matrix along the five morph levels in a given region, following a recent proposal^39^.

The estimated *R*^2^ from the nested models was used to assess general representations, as well as unique acoustic and categorical representations (Fig. 2f), as shown below:

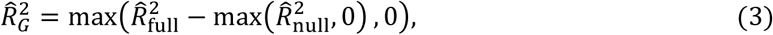

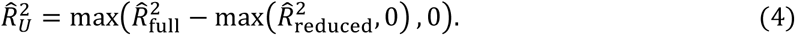

The max(·, 0) operator zeros any negative *R*^2^ that could result from NNLS fitting. While *R*^2^, in principle, cannot be negative, NNLS can yield a value below zero when the predictors explain less variance in the dependent variable than its mean, due to the non-negative weight constraint. In our case, this occurs particularly when the mean of distance estimates in a given neural RDM is negative. 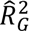 is a measure of general representations, estimating the variance explained by both model RDMs. 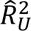 represents the unique variance explained by the acoustic or categorical RDM that is not included in the Reduced model. The nested model comparison, following Equation (4), enabled us to partition the variance explained by the highly correlated predictor RDMs 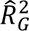, into their individual unique contributions (Supplementary Fig. 5, Supplementary Methods). The estimated 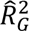 and 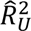 for each acoustic and categorial RDM were square-rooted and then Fisher-Z-transformed for visualisation and group-level inference, followed by baseline correction per region and participant to account for bogus effects driven by free parameters.

Additionally, time-resolved RSA was performed on surrogate data to validate the observed general representations of phonemes and prosody in each ROI (Supplementary Fig. 1). The surrogate data were created by permuting the corresponding five morph levels for each participant. All possible permutations of the five levels were considered, except for the original order (5! − 1 = 119). We measured the variance of a level-permuted neural RDM explained by both model RDMs at each time step, as described above, and took the median of the explained variance across permutations as the estimate of surrogate representations.

Lastly, we estimated the noise ceiling along the dimension of either phonemes or prosody in each ROI by calculating the lower and upper bounds of the structured variance in the corresponding neural RDM at each time step. The lower and upper bounds were computed for each participant following Equation (3), while the Full models were defined as follows:

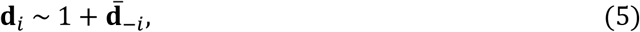

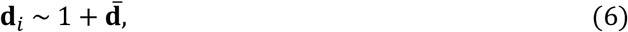

where ***d***_*i*_ represents the neural RDM for the *i*-th participant. The lower bound was estimated by predicting ***d***_*i*_ using the intercept and ***d***−_−*i*_, the neural RDM averaged across participants excluding the *i*-th participant as in Equation (5). The upper bound was derived by predicting ***d***_*i*_ using the intercept and ***d***−, the grand-averaged neural RDM as in Equation (6). The computed bounds were then averaged across participants, yielding a time-varying range of explainable variance in a given ROI (Supplementary Fig. 1).

### mTE analysis

To investigate how phonemic and prosodic information propagates between the involved ROIs, we applied mTE analysis to time-resolved neural RDMs. This analysis employed a recently proposed framework that combines Rényi’s *α*-order entropy, a generalisation of Shannon entropy, with a kernel trick to measure directed information transfer between two multivariate time series without estimating their individual or joint probability density functions^36,37^.

In this framework, the entropy of an arbitrary-dimensional time series **X** = {***x***_1_, ***x***_2_, ⋯, ***x***_T_} is measured as follows^36^:

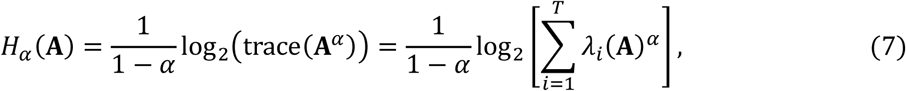

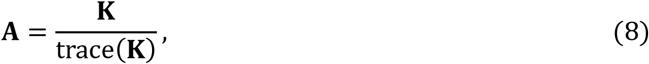

where **K** is the Gram matrix derived from the Gaussian kernel function *κ*(***x***_*i*_, ***x***_*j*_) = 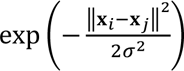. ***A*** is the normalised Gram matrix, and *λ* (***A***) is the *i*-th eigenvalue of ***A***. *H*_*α*_(***A***) is a matrix-based analogue to Rényi’s *α*-order entropy, which approaches Shannon entropy as *α* → 1. Given two multidimensional time series **X** and **X** of length **Y**, the matrix-based Rényi’s *α*-order joint entropy can be written as^36^:

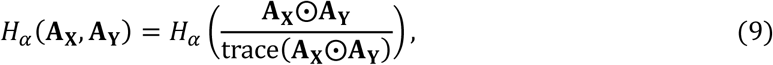

where ***A***_**X**_ and ***A***_**X**_ are the normalised Gram matrices by applying Equation (8) to the Gram matrices (**K**_**X**_)_*ij*_ = *κ*_**x**_(***x***_*i*_, ***x***_*j*_) and (**K**_**X**_)_*ij*_ = *κ*_**X**_(***y***_*i*_, ***y***_*j*_), respectively. The symbol ⨀ represents the Hadamard product. Based on Equations (7) and (9), mTE can be defined as^37^:

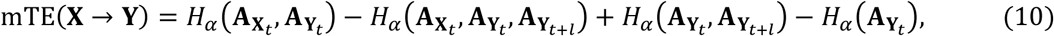

where ***A***_**X***t*_ and ***A***_**X***t*_ are the normalised Gram matrices of multivariate time series **X** and **X** at time *t*, and ***A***_**X***t*+*l*_ is the one based on **X** at future time *t* + *l*.

Using Equation (10), we assessed how informative the past RDMs in a source ROI **X**_*t*_ are for predicting the future RDMs in a target ROI **X**_*t*+*l*_ beyond its own past RDMs **X**_*t*_. This approach is similar to previous methods using Granger causality^40,41^ but can also capture non-linear interactions, including representational transformations from acoustics to categories. Here, we set the bandwidth of the Gaussian kernel function σ as the median distance between the time samples^70^, and used *α* = 2 for neutral weighting of rare events^36^. mTE analysis was performed on a subset of ROIs that showed significant general representations of either phonemes or prosody (Fig. 3), depending on the analysis of interest. For each of these ROIs, we first re-generated neural RDMs at each time sample from 0 to 800 ms after stimulus onset, without applying time windows, to avoid each sample from being informed by its past or future time points. Neural RDMs were built as described above in *Time-resolved RSA*, by computing neural pattern dissimilarities between all pairs of the five morph levels. These RDMs were vectorised and adaptively whitened, yielding a multivariate time series per ROI and participant, separately for phonemes and prosody. Next, we measured mTE between the time-resolved neural RDMs for pairs of ROIs, with each ROI alternatively serving as the source and target. The target RDM series was lagged by *l* to introduce time asymmetry, with the lag *l* varying from 4 to 300 ms to cover representational transfer across a broad time range. mTE was estimated at each time lag while ensuring that the number of time samples used remained equal.

Notably, the measured mTE values are non-negatively biased and differ in scales, as their upper bounds are determined by the entropy of the future RDMs in a target ROI conditioned on its own past: *H*_*α*_(***A***_**X**t+l_ |***A***_**X**t_) = *H*_*α*_(***A***_**X**t_, ***A***_**X**t+l_) − *H*_*α*_(***A***_**X**t_) in Equation (10)^42^.

These properties complicate the comparison of mTE values and statistical inference at the group level. Therefore, we normalised the mTE values by the mean and standard deviation of the surrogate mTE distribution, obtained by cyclically time-shifting the source RDM series based on a point exhaustively drawn from the interval between the 5^th^ and 95^th^ percentiles of the time samples used^43^. This procedure produced the normalised mTE (mTEn) value for each time lag, directed connection, and participant, separately for phonemes and prosody.

### Lateralisation analysis

Lateralisation analysis was performed to determine whether phonemic and prosodic representations at different abstraction levels were lateralised along ventral and dorsal stream regions. Specifically, we calculated the difference in the time series of baseline-corrected Fisher-Z variance uniquely explained by either acoustic or categorical RDM between homologous ROIs for each participant, separately for phonemes and prosody.

### Analysis of task-modulation effect

Here, we assessed whether phonemic and prosodic representations were enhanced by relevant task demands. First, we examined the task-modulation effect on the general representations of either phonemes or prosody within the subset of ROIs identified in the previous analysis (Fig. 3). For these ROIs, we ran time-resolved RSA on task-specific MEG source activity patterns to estimate the time-varying variance explained by both model RDMs during the phoneme and prosody identification tasks. We used the task-agnostic whitening matrix ***W*** for both tasks (see *Time-resolved RSA*), as there was no reason to assume a difference in the noise covariance structure across them. We then computed the difference between the tasks for each ROI and participant, separately for phonemic and prosodic representations.

Next, we examined the task-modulation effect at different abstraction levels, focusing on phonemic representations in the left pSTS, as no other effects were observed in the previous step. For each task, we estimated the time-varying unique variance explained by each acoustic and categorical RDM using variance partitioning. We then calculated the difference between the tasks for each model RDM and participant.

Finally, we tested the lateralisation of the task-modulation effect on the categorical representations of phonemes in the left pSTS, as no effect was observed for the acoustic representations. We calculated the difference between the tasks for the categorical RDM in the right pSTS and contrasted this difference with that in the left pSTS for each participant.

The (unique) variance explained, measured in the analysis, was converted to the Fisher-Z scale and baseline-corrected before being compared between the tasks.

### Statistical inference

For group-level inference in all MEG data analyses, statistical significance was determined using cluster-based permutation tests. Specifically, either temporal or spatiotemporal clusters were formed by one-sided *t*-tests against zero with a threshold of *p* < 0.05, and family-wise error rate was controlled at *p* < 0.05 across time points per ROI, across time lags per directed connection in the mTE analysis, or across time points and vertices in the searchlight analysis. To assess the difference between the two tasks, we performed one-sided instead of two-sided *t*-tests as we anticipated that task demands would enhance relevant representations. Two-sided *t*-tests were used to compare homologous ROIs when testing for lateralisation. Permutation distributions were generated by randomly shuffling signs or condition labels 100,000 times in the ROI-based, and 10,000 times in the searchlight analyses. Comparisons across multiple ROIs or directed connections were controlled at a false discovery rate of *q* < 0.05, based on the minimum cluster *p*-value for each contrast.

## Data availability

The stimuli presented and the behaviour responses of individual participants collected during the MEG experiment, as well as the preprocessed neural data—including the neural RDMs and time-resolved RSA and mTE results based on the ROIs—are publicly available and can be downloaded via the following repository: https://github.com/SeungCheolBaek/representation_dynamics_phonemes_prosody.

## Code availability

The code to replicate the ROI-based RSA and mTE analysis, as well as the main figures, is publicly available in the following repository: https://github.com/SeungCheolBaek/representation_dynamics_phonemes_prosody.

## Acknowledgements

We thank Florian Scharf for extensive efforts in stimulus generation, and express our gratitude to Yvonne Wolff-Rosier for assistance in conducting the experiment and collecting the data. This work was supported by the Max Planck Society. The funder had no role in study design, data collection and analysis, decision to publish or preparation of the manuscript.

## Author contributions

Conceptualization: S-C.B., M.G., and D.S. Methodology: S-C.B., S-G.K., M.G., and D.S. Software: S-C.B., B.M., M.G., and D.S. Validation: S-C.B., S-G.K., B.M., M.G., and D.S. Formal analysis: S-C.B., M.G., and D.S. Data curation: S-C.B., B.M., M.G., and D.S. Writing–original draft: S-C.B. and D.S. Writing–review and editing: S-C.B., S-G.K., B.M., M.G., and D.S. Visualization: S-C.B., S-G.K., and D.S. Supervision: D.S. Project administration: D.S. Funding acquisition: D.S. All authors have read, commented on and approved the manuscript.

## Competing interests

The authors declare no competing interests.

## Additional information

**Correspondence and requests for materials** should be addressed to Daniela Sammler.

## Notes

https://github.com/SeungCheolBaek/representation_dynamics_phonemes_prosody

### Competing Interest Statement

The authors have declared no competing interest.

### Summary of Updates

Substantial changes in the Introduction and the Discussion to clarify the research questions and theoretical contributions of this study; Supplementary information updated.

