## Supplementary Information for "Dynamic acoustic-to-categorical representations of phonemes and prosody along ventral and dorsal speech streams"

### Supplementary Methods

#### Behavioural pre-tests

During the magnetoencephalography (MEG) experiment, participants listened to a set of 25 single words per speaker, gradually varying in word-initial phonemes (from "Bar" to "Paar") and prosody (from statement to question). The stimulus set was tailored individually for each participant by selecting 5 x 5 levels from the 61 x 61-step orthogonal morphing continua of voice onset time (VOT) and pitch contour of each speaker (middle panel of Fig. 1a). The selection was based on each participant's performance in two behavioural pre-tests conducted on a separate day before the MEG experiment, using Presentation® software (version 18.0, Neurobehavioral Systems). The two pre-tests lasted about two hours, including breaks in between.

**Pre-test 1.** *Procedure*. In the first pre-test, we aimed to determine the position and sharpness of the individual perceptual boundaries of two phoneme and prosody categories, to select five stimulus levels around the category boundaries that ensured matched difficulty levels in the two identification tasks. To achieve this, we employed an adaptive-staircase method[1], separately for VOT (phoneme) and pitch contour (prosody).

Pre-test 1 consisted of three repeated sessions. Each session contained two phoneme and two prosody blocks, one of each for the male and the female speaker. In these blocks, participants were asked to identify either the word ("Bar"/"Paar," phoneme block) or the prosody of the stimuli ("statement"/"question," prosody block). Phoneme and prosody blocks of the same speaker immediately followed each other. The respective task was indicated at the beginning of each block. During the block, the colour of a fixation cross in the centre of the screen reminded participants of the task type (green for phonemes and red for prosody). The two corresponding response options were constantly shown on the left and right side of the screen. The order of task and speaker was randomised across sessions and participants. Before the first pre-test proper, participants had a mini-session to familiarise themselves with the tasks and to adjust the sound intensity to a comfortable level. Pre-test 1 lasted about an hour, with optional breaks between sessions.

In each block, we adaptively tracked participants' categorical percepts in response to various morph steps along the task-relevant dimension using two streams. These two tracking streams started at different positions on the target dimension (9$^{th}$ and 29$^{th}$ morph steps for phonemes, and 11$^{th}$ and 41$^{st}$ for prosody), thus taking different categories as their references. We interleaved the two streams pseudorandomly to prevent participants from knowing which stream they were following, such that stimuli from the same stream were not repeated more than three times in a row. Each stream consisted of up to 50 trials, in each of which participants reported their percept to a given stimulus by pressing a button on a response box corresponding to the options displayed on the screen. Response button assignments to the two phoneme or prosody categories was counterbalanced between odd- and even-numbered participants. Responses were accepted until 1500 ms after stimulus onset, at which point the next trial began.

For each stream, if the current response matched its reference category, the morph step for the next trial was adjusted by one hop (one-up) towards the opposite extreme from its starting position (hop size: phonemes $= 2$, prosody $= 4$ morph steps). Otherwise, the morph step was adjusted by two hops (two-down) towards the extreme on the same side. At the same time, the task-irrelevant dimension varied randomly across trials based on a predefined pool (11 morph steps, phonemes ranged from 11$^{th}$ to 31$^{th}$ steps in 2-step-sized intervals; prosody ranged from 1$^{st}$ to 51$^{st}$ steps in 5-step-sized intervals). During this adaptive tracking, we monitored reversals in responses, where the current response differed from the previous one within a given stream. Each block continued until either 12 reversals were identified or 50 trials were presented, for both tracking streams.

To help participants swiftly adapt to a given task, six practice trials were provided at the beginning of each block. These trials consisted of stimuli with morph steps close to the two ends of the task-relevant dimension and were not considered in the analysis.

*Analysis and level selection*. Participants' behavioural responses were analysed in MATLAB (R2014a, MathWorks). For each participant and session, we first fitted a psychometric function (@PAL_Logistic) to the behavioural responses for each task and speaker using the *PAL_PFML_Fit* function in the Palamedes toolbox[2] (version 1.7.0). *PAL_PFML_Fit* searched for the 50%-point boundary $a$ (in morph step) and slope $b$ of a psychometric function to maximise their likelihood based the observed proportions of "Paar" and "question" responses at the given morph steps (guess rate and lapse rate parameters were fixed at 0.01). The fitted psychometric function was then used to identify outliers within each session, defined as the observed proportion that deviated from the expected proportion by more than 0.33. After rejecting the outliers, we again fitted a psychometric function as described above, this time to the behavioural data aggregated across the three sessions. This yielded single estimates of the 50%-point boundary $a$ and slope $b$ for each task, speaker, and participants.

Next, the estimated parameters were used to select five individualised levels of phonemes and prosody. Specifically, we selected five morph steps from the 61-step phoneme or prosody continuum that were closest to $a$, $a \pm \frac{3}{2b}$, and $a \pm \frac{3}{b}$ based on the corresponding parameters. The selected morph steps for phonemes and prosody were combined to generate an individualised set of 25 distinct stimuli (i.e., 5 x 5 levels) for each speaker (see also left panel in Fig. 2d). These stimuli were presented in the second pre-test.

**Pre-test 2.** *Procedure*. The second pre-test was to validate the stimulus set created for each participant in the first pre-test. Pre-test 2 was conducted after a short break following Pre-test 1, and also consisted of three sessions, each of which contained 4 blocks (two phoneme, two prosody blocks, one of each for each speaker). Participants listened to the identical set of stimuli in the same order over the two consecutive blocks of the same speaker. Counterbalancing of speakers and tasks was as in Pre-test 1. Again, the task was indicated at the beginning of each block, with the colour of the fixation cross (green for phonemes and red for prosody) and the two corresponding alternative options on the screen serving as reminders throughout the block. Pre-test 2 also lasted about an hour, with optional breaks between sessions.

Each block consisted of 56 trials, with the first six trials serving as practice and excluded from the analysis. Across the remaining 50 trials, 25 distinct stimuli were presented in a pseudorandomised order following the principles of a type-1 index-1 sequence to prevent carry-over and position effect[3]. In each trial, participants listened to a single word and classified the stimulus category according to the task by pressing a button on a response box. Button assignment was counterbalanced across participants. Responses were accepted until 2500 ms after stimulus onset, followed by the stimulus of the next trial.

*Analysis and level adjustment*. Similar to Pre-test 1, participants' behavioural responses were analysed using the Palamedes toolbox[2] and custom code in MATLAB. For each participant, we fitted a psychometric function (@PAL_Logistic) to behavioural responses aggregated across the three sessions using the *PAL_PFML_Fit* function (guess rate and lapse rate parameters were fixed at 0.01). This yielded single estimates of the 50%-point boundary $a$ and slope $b$ for each task, speaker, and participants.

We visually inspected the actual behavioural responses and the fitted psychometric function. If necessary, we adjusted the five individualised levels for phonemes and prosody to better align with $a$, $a \pm \frac{3}{2b}$, and $a \pm \frac{3}{b}$, derived from the newly estimated parameters. Then, we combined the adjusted five levels of phonemes and prosody to generate an individualised set of 25 distinct stimuli per speaker for the MEG experiment.

#### Estimation of the pitch divergence point

In our manipulation of the experimental stimuli, phonemes were characterised by a relatively clearly defined acoustic cue, namely the timing of voice onset. In contrast, prosodic contours were characterised by the continuous change in the fundamental frequency ($F_0$) of the words over time, making it difficult to define a clear acoustic landmark to which neural and behavioural responses were time-locked. To still obtain a proxy of such a landmark, we estimated the point in the middle of the word where the pitch contours of the five prosody levels clearly diverged from one another (Supplementary Fig. 3).

To estimate the pitch divergence point, we first extracted the pitch contours from all the stimuli presented during the MEG experiment. The pitch extraction was performed using the *To Pitch (cc)* function in PRAAT[4] (version 6.3.14), which computed time-varying $F_0$ values based on cross-correlation. The $F_0$ values were estimated every 1 ms for each stimulus, with the pitch floor parameter set to 60 Hz for the male speaker and 75 Hz for the female speaker. All other parameters were set to PRAAT's default values. Next, we determined the intersection points for pitch contour pairs of the five prosody levels, for each phoneme level, speaker, and participant. Specifically, we computed the differences in time-varying $F_0$ values and identified the zero-crossing points (i.e., where the sign of the difference values changed). Notably, the zero-crossing points were dispersed over time for different pitch contour pairs and were also observed at the temporal edges due to noise in pitch extraction. To address this, we fitted a one-dimensional Gaussian mixture model (2-3 components) to the zero-crossing points identified across all pairs of the five prosody levels using the *GaussianMixture* function in the scikit-learn toolbox (version 1.1.2). We considered the mean of a Gaussian component between 100-400 ms after stimulus onset as the pitch divergence point for a given phoneme level, speaker, and participant. As shown by the grey rectangles in Supplementary Fig. 3, the pitch divergence point ranged from 184-214 ms post-onset for the male speaker and from 221-250 ms post-onset for the female speaker. In Figs. 3 and 5, the range of the pitch divergence points was aggregated across speakers.

#### Correlation between acoustic and categorical representational dissimilarity matrices (RDMs)

In the time-resolved representational similarity analysis (RSA), we built two model RDMs with respect to phonemes and prosody to explore their acoustic and categorical representations in the brain. Acoustic RDMs were based on the original morph steps of the stimuli presented to each individual participant along the 61-step phoneme or prosody continuum, while categorical RDMs were derived from the expected proportions of “Paar” or “question” responses at the five morph levels, estimated from psychometric curves fitted to each participant’s behaviour (Fig. 2d,e). However, because both the original morph steps and expected proportions increased monotonically across the five phoneme or prosody levels, this led to a correlation between acoustic and categorical RDMs. The multicollinearity induced by such correlations can make it challenging to disentangle the contribution of the respective model RDMs to explaining the variance of the neural RDMs at specific regions and time points.

To estimate the strength of the correlation between acoustic and categorical RDMs, we calculated Pearson’s correlation based on their lower triangular entries (as RDMs are diagonally symmetric). The correlation was measured for each participant, separately for phonemes and prosody. As shown in Supplementary Fig. 5, acoustic and categorical RDMs were highly correlated, both for phonemes (mean correlation ± the standard error of the mean (SEM) $= 0.915 \pm 0.016$; two-sided inference against zero after Fisher-Z transformation, $t_{28} = 17.263$, $p = 1.853E-16$, Cohen’s d $= 3.206$) and prosody (mean ± SEM $= 0.903 \pm 0.010$; two-sided inference against zero after Fisher-Z transformation, $t_{28} = 21.527$, $p = 5.842E-19$, Cohen’s d $= 3.997$). These high correlations called for variance partitioning based on nested linear modelling to tease apart the unique contributions of the two model RDMs, which allowed us to explore acoustic and categorical representations of phonemes and prosody (see *Time-resolved RSA* in Methods).

#### Validation of modelling stimulus acoustics

To build the acoustic RDMs, we modelled stimulus acoustics as the original morph steps of the stimuli along the 61-step phoneme or prosody continuum. We considered this approach based on parameterised morphing more robust than the use of hand-crafted features, which may produce variable outcomes depending on the number and types of features considered and the variability in feature extraction (e.g., hyperparameters). However, before doing so, we validated that the morph steps corresponded to respective phonemic and prosodic cues using 26 hand-crafted acoustic features. These acoustic features included eight time-related features (duration, minimum $F_0$ time, maximum $F_0$ time, onset $F_0$ time, offset $F_0$ time, middle $F_0$ time, minimum intensity time, and maximum intensity time in seconds), 13 frequency-related features (minimum $F_0$, maximum $F_0$, range of $F_0$, onset $F_0$, offset $F_0$, middle $F_0$, slope of $F_0$ throughout the stimulus, slope of $F_0$ for the first half, slope of $F_0$ for the second half, mean $F_0$, median $F_0$, and standard deviation of $F_0$ in Hertz, and slope of $F_0$ in semitones per second), and five intensity-related features (minimum intensity, maximum intensity, maximum difference in intensity, mean intensity, standard deviation of intensity in decibels). We extracted these acoustic features from the stimuli presented during the MEG experiment using PRAAT[4] (see *Estimation of the pitch divergence point* in Supplementary Methods for details on pitch extraction) and, if needed, manually corrected the estimation of onset $F_0$ time (as it corresponds to our manipulation of VOT for phonemes). The statistics of the extracted acoustic features are summarised in Supplementary Table 1.

To examine which acoustics are represented in the original morph steps, we performed principal component analysis (PCA) on the 26 extracted acoustic features, separately for the stimuli of the male and the female speaker. We first standardised each feature to have zero mean and unit variance and estimated the principal components (PCs) of the feature set, such that 99% of the total variance was explained. Then, we mapped the individual stimuli, coded with the original morph steps along the two-dimensional phoneme and prosody continua, onto the two largest PCs (each explaining 51.45% and 27.62% of the total variance for the male, and 51.24% and 26.80% for the female speaker). As shown in Supplementary Fig. 9a, the first and second PCs were respectively well-aligned with the gradient of phoneme and prosody morph steps for both the male and the female speaker, confirming that the original morph steps represent stimulus acoustics.

For the quantitative evaluation of this mapping, we computed the correlation between the original morph steps along either the phoneme or prosody continuum and the PC scores of the stimuli (Supplementary Fig. 9b). The scores from the first PC showed an almost perfect correlation with the prosody morph steps (male: $r = 0.975$, 95% bootstrapping confidence interval (CI) = $[0.973, 0.978]$, 1000 resamples with replacement; female: $r = 0.995$, 95% bootstrapping CI = $[0.994, 0.995]$; 1000 resamples with replacement), but low or no correlation with the phoneme morph steps. In contrast, the scores from the second PC displayed the opposite pattern (correlation with the phoneme morph steps, male: $r = 0.962$, 95% bootstrapping CI = $[0.958, 0.967]$, 1000 resamples with replacement; female: $r = 0.980$, 95% bootstrapping CI = $[0.977, 0.982]$, 1000 resamples with replacement). For the other PCs, the scores showed low or almost no correlations with either the phoneme or prosody morph steps. These results, again, suggest that the acoustic features are well-reflected in the original morph steps, licensing their use for modelling stimulus acoustics.

### Supplementary Figures

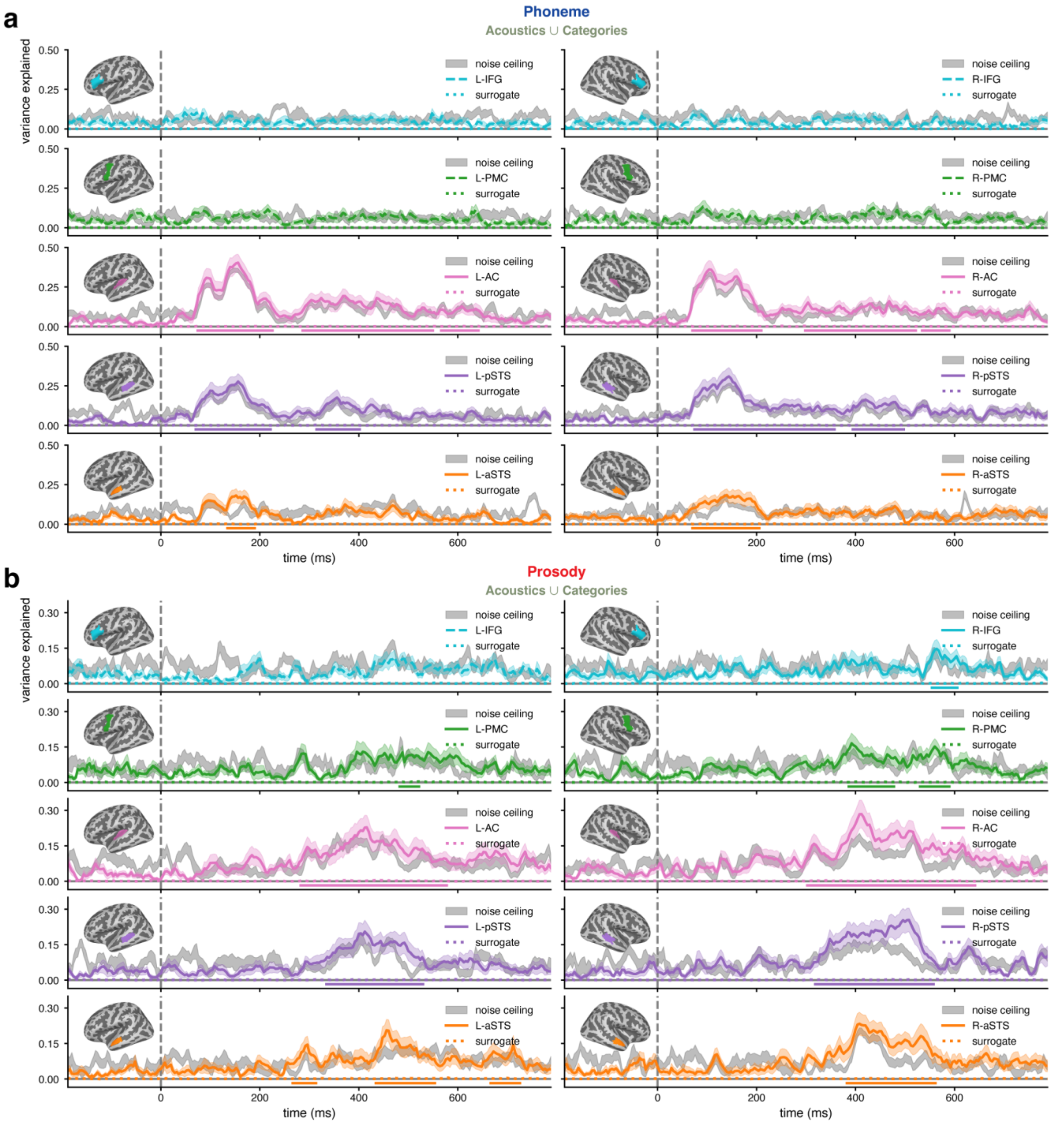

**Supplementary Fig. 1| Phonemic and prosodic representations in regions of interest (ROIs). a**,**b**, Time-varying variance explained by both acoustic and categorical representational dissimilarity matrices (RDMs) for phonemes (**a**) and prosody (**b**) in the left- (L-; left column) and right-hemispheric (R-; right column) ROIs. Solid lines indicate ROIs with significant representations; otherwise, dashed lines are used. Shaded areas denote the standard error of the mean (SEM). Horizontal lines above the x-axis show significant time intervals. Grey areas represent empirical noise ceiling. Surrogate representations are shown as dotted lines. IFG, inferior frontal gyrus; PMC, premotor cortex; AC, auditory cortex; p/aSTS, posterior/anterior superior temporal sulcus.

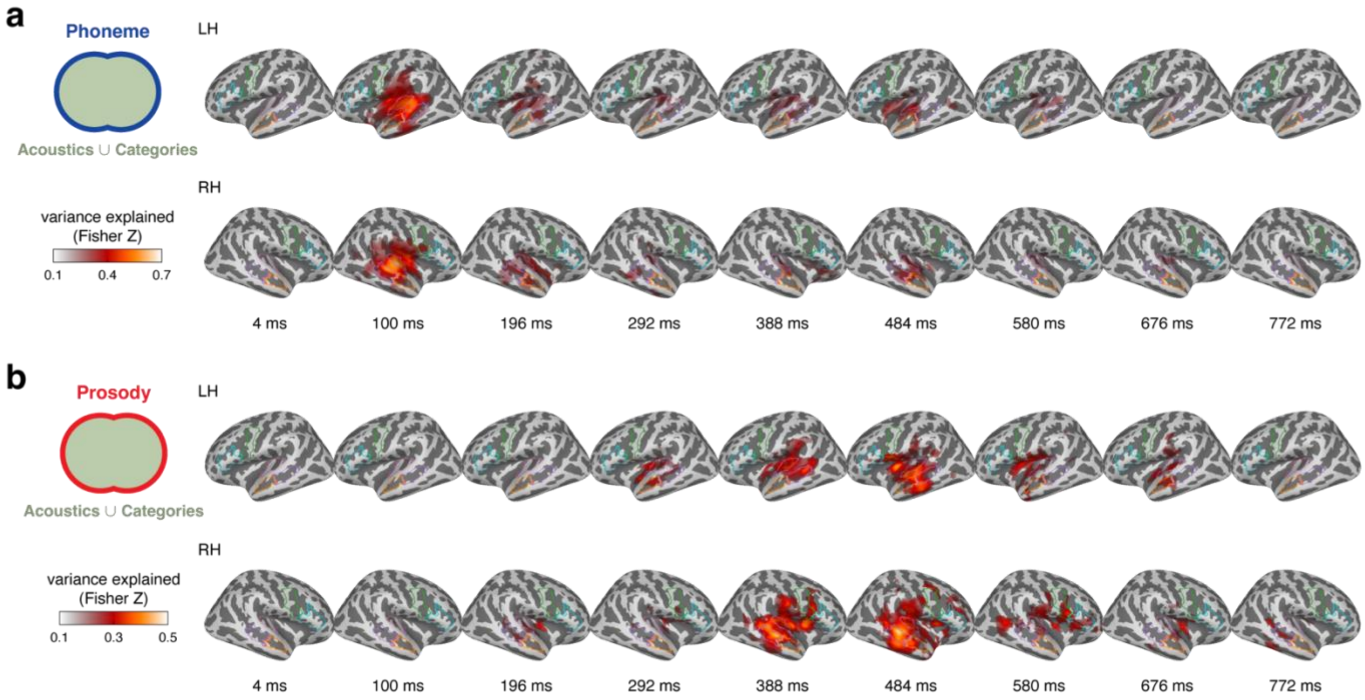

**Supplementary Fig. 2| Cortical representations of phonemes and prosody from the whole-brain searchlight analysis. a,b**, Cortical variance map series (Fisher-Z scale, baseline-corrected) of significant phonemic (**a**) and prosodic (**b**) representations in the left (LH; upper) and right (RH; lower) hemispheres. The spatial extents of the ROIs are outlined in different colours (see Fig. 2a) for comparison with Fig. 3.

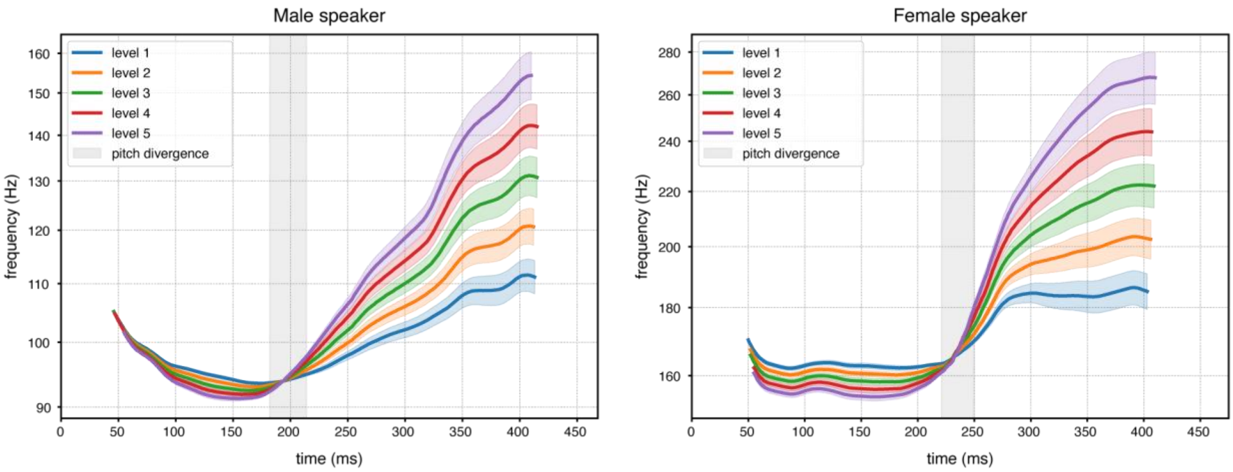

**Supplementary Fig. 3| Pitch contours of the stimuli and the range of pitch divergence points.** Pitch contours of the stimuli across the five prosody levels for the male (left panel) and female (right panel) speaker. For each prosody level, pitch contours were first averaged over the five phoneme levels and then across participants (thick lines). Shaded areas indicate SEM. Grey rectangles represent the range of the pitch divergence points.

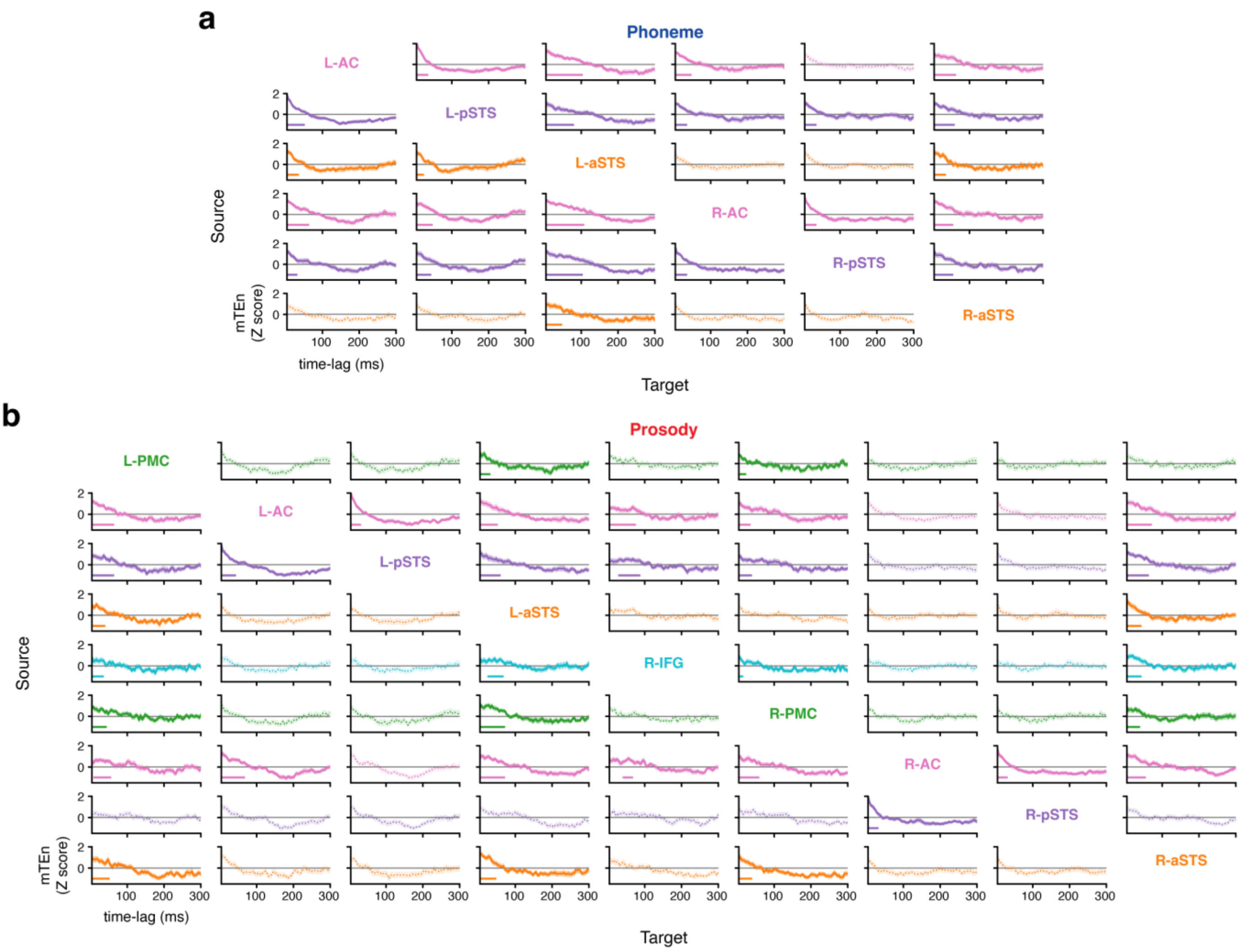

**Supplementary Fig. 4| Transfer of phonemic and prosodic representations between the involved ROIs across time lags. a,b**, normalised multivariate transfer entropy (mTEn) values from the source ROIs (in rows) to the target ROIs (in columns) across time lags for phonemes (**a**) and prosody (**b**). Directed connections are colour-coded according to the source ROIs. Solid lines indicate directed connections with significant representational transfer; otherwise, dotted lines are used. Shaded areas denote SEM. Horizontal lines above the x-axis show significant time lag intervals. L-, left; R-, right; IFG, inferior frontal gyrus; PMC, premotor cortex; AC, auditory cortex; p/aSTS, posterior/anterior superior temporal sulcus.

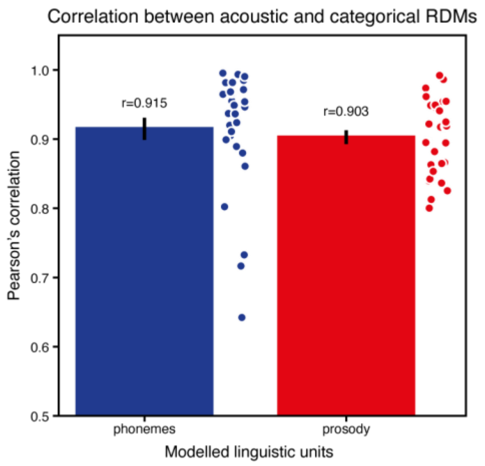

**Supplementary Fig. 5| Correlation between acoustic and categorical representational dissimilarity matrices (RDMs).** Pearson's correlations were calculated between the lower triangular entries of acoustic and categorical RDMs, separately for phonemes (blue) and prosody (red). Error bars indicate SEM. Circles denote individual participants.

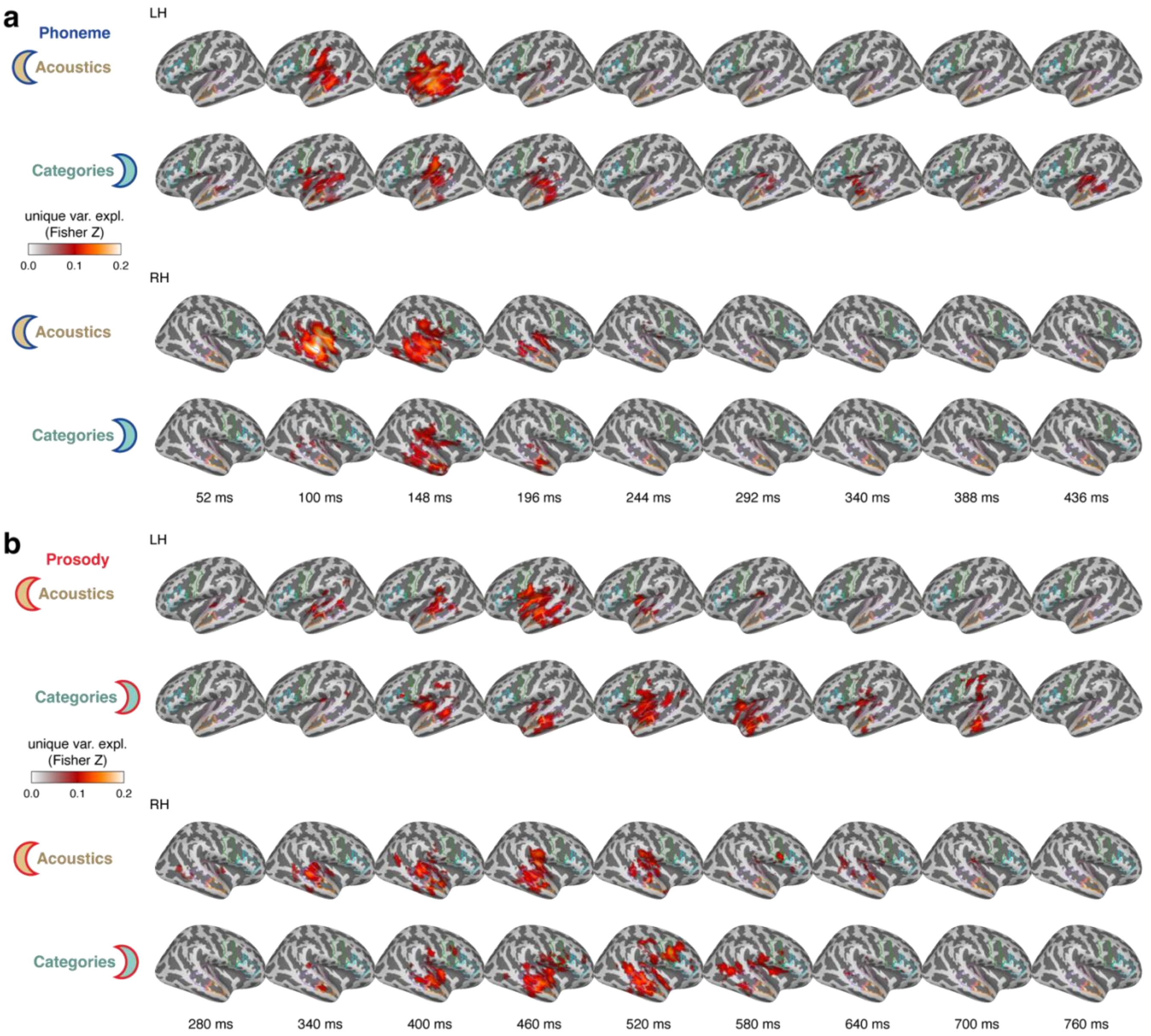

**Supplementary Fig.6| Cortical representations of acoustic and categorical information for phonemes and prosody from the whole-brain searchlight analysis.** Cortical variance map series (Fisher-Z scale, baseline-corrected) of significant unique representations (cluster-based permutation tests, one-sided $t_{28}$-test against zero, cluster-forming threshold at $p < 0.05$, 10,000 permutations, family-wise error rate $= 0.05$ across vertices and time points) for phonemes (**a**) and prosody (**b**) in the left (LH; upper panel) and right (RH; lower panel) hemispheres. Acoustic representations are shown in the first row, and categorical representations in the second row. The spatial extents of the ROIs are outlined in the corresponding colours (see Fig. 2a) for comparison with Fig. 5.

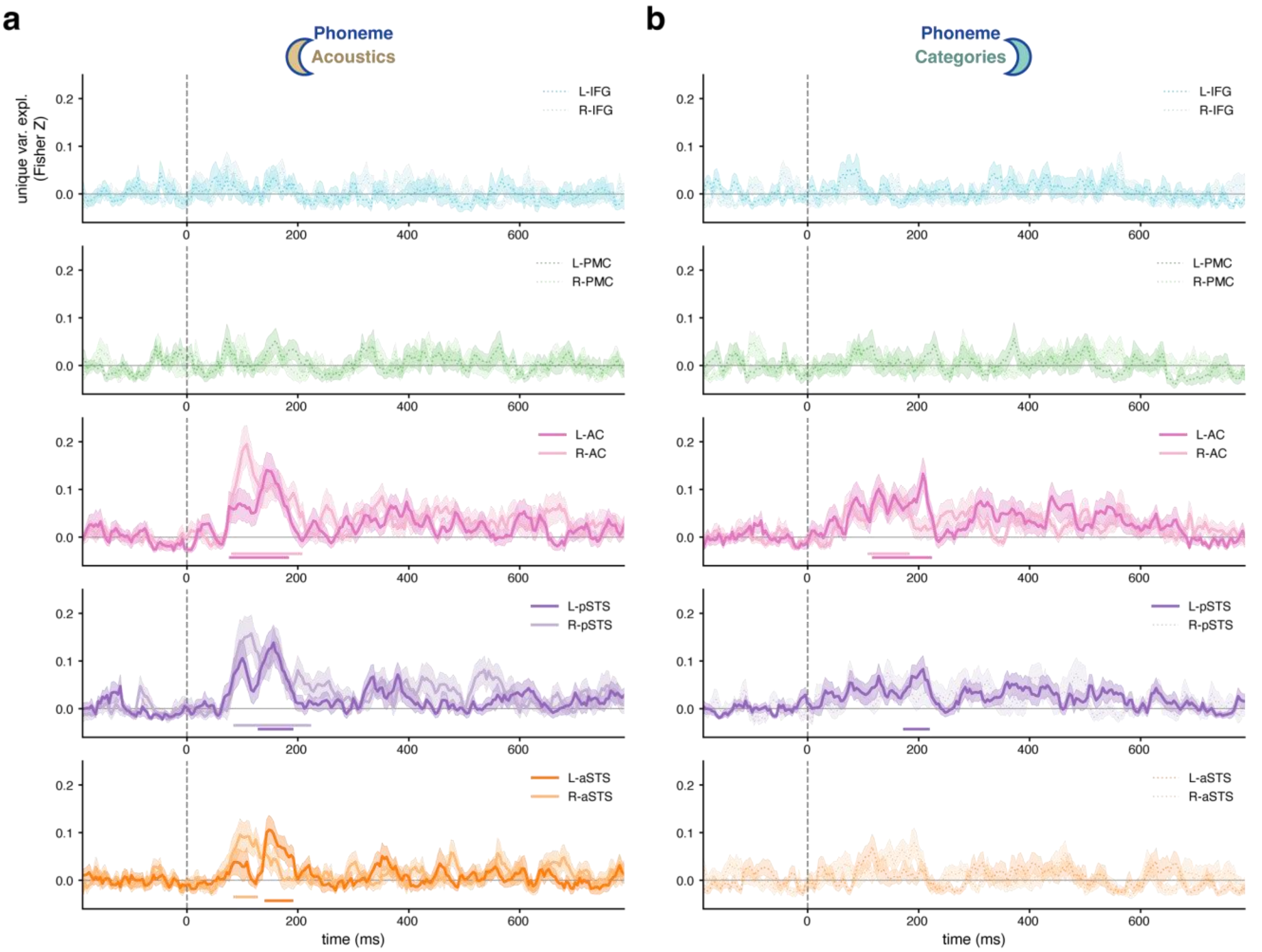

**Supplementary Fig. 7| Comparison of acoustic and categorical representations for phonemes between homologous ROIs. a,b**, Time-varying variance (Fisher-Z scale, baseline-corrected) uniquely explained by either acoustic (**a**) or categorical (**b**) RDM for phonemes in all ROIs. Homologous ROI pairs are grouped in rows, with right-hemispheric ROIs shown in pale colour. Solid lines indicate ROIs with significant unique representations (significant time intervals are shown as horizontal lines above the x-axis); otherwise, dotted lines are used. Shaded areas denote SEM. Neither acoustic (**a**) nor categorical (**b**) representations were lateralised in any homologous ROI pairs (cluster-based permutation tests, two-sided $t_{28}$-test, all cluster $p > 0.05$). L-, left; R-, right; IFG, inferior frontal gyrus; PMC, premotor cortex; AC, auditory cortex; p/aSTS, posterior/anterior superior temporal sulcus.

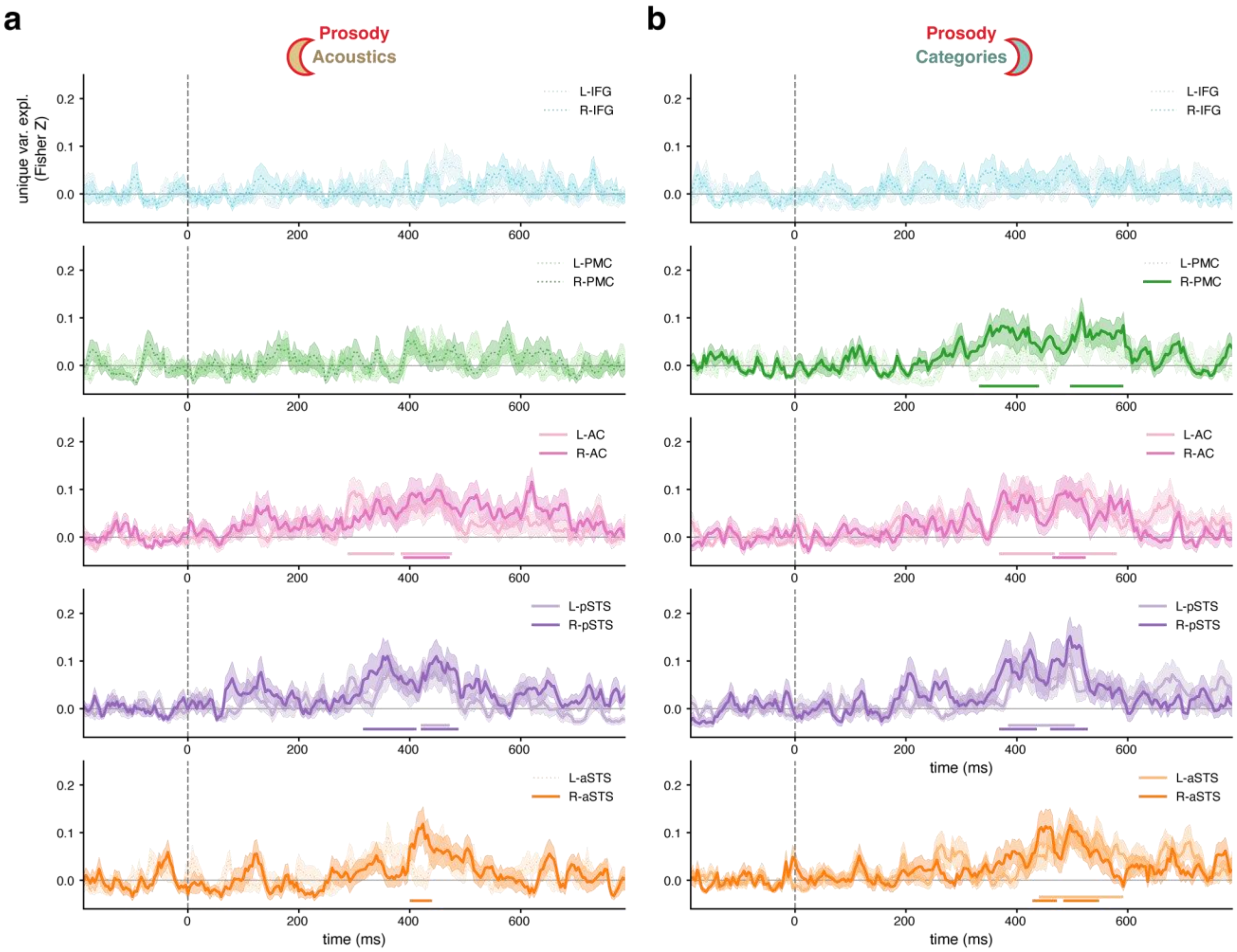

**Supplementary Fig. 8| Comparison of acoustic and categorical representations for prosody between homologous ROIs. a,b**, Time-varying variance (Fisher-Z scale, baseline-corrected) uniquely explained by either acoustic (**a**) or categorical (**b**) RDM for prosody in all ROIs. Homologous ROI pairs are grouped in rows, with left-hemispheric ROIs shown in pale colour. Solid lines indicate ROIs with significant unique representations (significant time intervals are shown as horizontal lines above the x-axis); otherwise, dotted lines are used. Shaded areas denote SEM. Neither acoustic (**a**) nor categorical (**b**) representations were lateralised in any homologous ROI pairs (cluster-based permutation tests, two-sided $t_{28}$-test, all cluster $p > 0.05$). L-, left; R-, right; IFG, inferior frontal gyrus; PMC, premotor cortex; AC, auditory cortex; p/aSTS, posterior/anterior superior temporal sulcus.

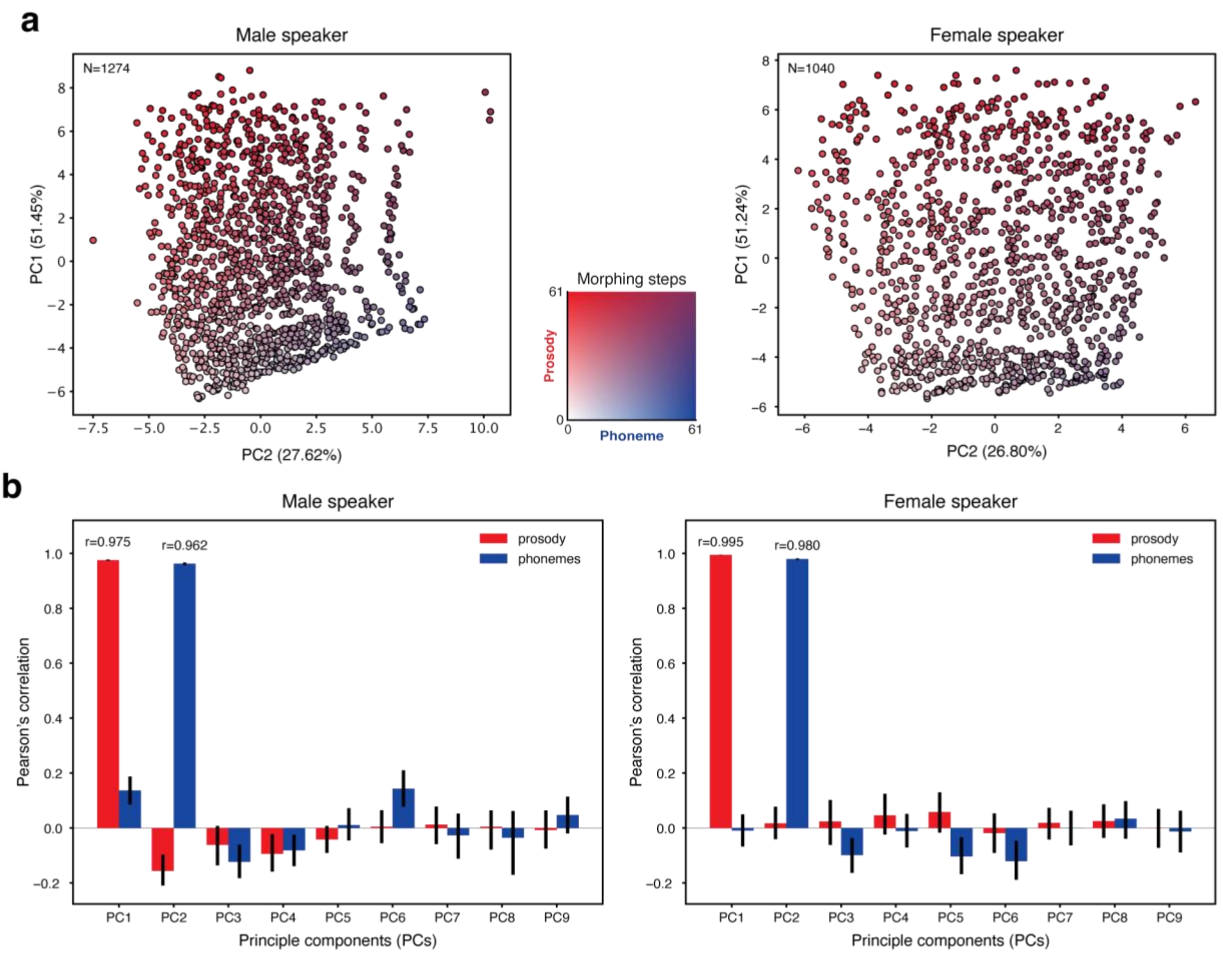

**Supplementary Fig. 9| Comparison of the original morph steps and the principal components (PCs) of the acoustic feature set. a**, Mapping of the stimuli, coded with the original morph steps along the two-dimensional orthogonal continua of phonemes and prosody (inset in the middle), onto the two strongest PCs from the set of 26 hand-crafted acoustic features for the male (left panel) and the female (right panel) speaker. Dots represent individual stimuli, colour-coded according to the inset in the middle. **b**, Correlation between the original morph steps, with respect to phonemes (blue) and prosody (red), and the scores of the stimuli from the first to the ninth PCs (together explaining >99% of the total variance) for the male (left panel) and the female (right panel) speaker. Error bars indicate 95% bootstrapping (1000 resamples with replacement) confidence intervals.

### Supplementary Tables

**Supplementary Table 1| Summary statistics of stimulus acoustic features.**

| Voice | Phoneme | | Prosody | | Duration (s) |
|---|---|---|---|---|---|
| | Nsteps | step range | Nsteps | step range | |
| male | 26 | 4-38 | 49 | 8-60 | 0.478±0.009 |
| female | 26 | 4-31 | 40 | 14-60 | 0.479±0.009 |

| | **Tmin$F_0$ (s)** | **Tmax$F_0$ (s)** | **Ton$F_0$ (s)** | **Toff$F_0$ (s)** | **Tmiddle$F_0$ (s)** |
|---|---|---|---|---|---|
| male | 0.205±0.079 | 0.319±0.176 | 0.040±0.009 | 0.448±0.011 | 0.245±0.010 |
| female | 0.197±0.088 | 0.323±0.166 | 0.045±0.009 | 0.441±0.009 | 0.243±0.009 |

| | **TminInt (s)** | **TmaxInt (s)** | **min$F_0$ (Hz)** | **max$F_0$ (Hz)** | **range$F_0$ (Hz)** |
|---|---|---|---|---|---|
| male | 0.443±0.010 | 0.063±0.016 | 91.40±2.57 | 138.20±27.69 | 46.81±29.02 |
| female | 0.436±0.009 | 0.145±0.076 | 156.35±4.44 | 237.05±48.66 | 80.70±52.65 |

| | **on$F_0$ (Hz)** | **off$F_0$ (Hz)** | **middle$F_0$ (Hz)** | **gradientAll (Hz)** | **gradient1st (Hz)** |
|---|---|---|---|---|---|
| male | 111.60±5.48 | 132.63±29.74 | 98.28±3.73 | 21.07±28.75 | -13.32±6.13 |
| female | 186.93±4.87 | 229.74±55.36 | 162.59±1.50 | 42.81±55.77 | -24.33±5.40 |

| | **gradient2nd (Hz)** | **mean$F_0$ (Hz)** | **median$F_0$ (Hz)** | **stdev$F_0$ (Hz)** | **Slope$F_0$ (st/s)** |
|---|---|---|---|---|---|
| male | 34.39±26.18 | 107.18±8.79 | 102.35±4.98 | 14.78±10.25 | 27.20±11.15 |
| female | 67.15±55.74 | 185.47±14.86 | 170.70±3.02 | 29.92±21.53 | 31.97±12.16 |

| | **minInt (dB)** | **maxInt (dB)** | **maxDiffInt (dB)** | **meanInt (dB)** | **stdevInt (dB)** |
|---|---|---|---|---|---|
| male | 48.28±1.83 | 71.93±0.77 | 23.64±2.33 | 64.70±0.78 | 5.94±0.90 |
| female | 49.45±1.82 | 69.96±0.50 | 20.51±2.01 | 65.82±0.53 | 6.04±0.98 |

This table lists mean ± 1 standard deviation for each of 26 acoustic features extracted from the stimuli for male and female voices, alongside the number of morph steps (Nsteps) selected and their range (step range) across participants out of the 61-step phoneme or prosody continuum. Tmin$F_0$, minimum $F_0$ time; Tmax$F_0$, maximum $F_0$ time; Ton$F_0$, onset $F_0$ time; Toff$F_0$, offset $F_0$ time; Tmiddle$F_0$, middle $F_0$ time; TminInt, minimum intensity time; TmaxInt, maximum intensity time; min$F_0$, minimum $F_0$; max$F_0$, maximum $F_0$; range$F_0$, range of $F_0$; on$F_0$, onset $F_0$; off$F_0$, offset $F_0$, middle$F_0$, middle $F_0$; gradientAll, slope of $F_0$ throughout the stimulus; gradient1st, slope of $F_0$ for the first half; gradient2nd, slope of $F_0$ for the second half; mean$F_0$, mean $F_0$; median$F_0$, median $F_0$; stdev$F_0$, standard deviation of $F_0$; slope$F_0$, slope of $F_0$ in semitones per second; minInt, minimum intensity; maxInt, maximum intensity; maxDiffInt, maximum difference in intensity; meanInt, mean intensity; stdevInt, standard deviation of intensity.
